# Identification of DksA as a novel pro-inflammatory mediator of *Pseudomonas aeruginosa* under conditions mimicking chronic cystic fibrosis lung infection

**DOI:** 10.64898/2025.12.21.695867

**Authors:** Merel Wauters, Laura Bollé, Sara Van den Bossche, Lucia Grassi, Delphi Van Haver, Sara Dufour, Simon Devos, Francis Impens, Eva Van Braeckel, Anna K.H. Hirsch, Marvin Whiteley, Xavier Saelens, Aurélie Crabbé

**Author notes:** ***Corresponding author*** Aurélie Crabbé.

## Abstract

Chronic infection with *Pseudomonas aeruginosa* is a major driver of airway inflammation, which plays a central role in the progression of cystic fibrosis (CF) lung disease. During long-term colonization, *P. aeruginosa* adapts to the CF lung by downregulating virulence factors and adopting a biofilm-associated, mucoid lifestyle. Despite the expected reduction in immune activation due to these adaptations, excessive inflammation persists, a paradox that remains poorly understood.

Our objective was to identify novel bacterial mediators sustaining persistent inflammation by *P. aeruginosa* in the CF lung. To this end, we analyzed clinical *P. aeruginosa* CF isolates, cultured them in synthetic CF sputum medium, and exposed 3D lung epithelial cell cultures to the resulting cell-free supernatants. There was considerable variability in pro-inflammatory activity among the isolates, with a subset of the isolates inducing strong IL-8 secretion by the 3D cells despite low production of known virulence factors. Comparative proteomics analysis of the cell-free supernatants of pro- and anti-inflammatory isolates revealed several mediators not previously linked to inflammation.

Thirteen of these candidate pro-inflammatory mediators were selected for further analysis. Using *P. aeruginosa* transposon mutants lacking the respective mediators, DksA (a transcription factor) was confirmed as an immunomodulatory mediator in the 3D lung model. Finally, analysis of existing transcriptomes of *P. aeruginosa* in CF sputum, revealed that *dksA* was found to be one of the most strongly expressed genes in this patient population, highlighting the relevance of our findings. In conclusion, we identified a novel *P. aeruginosa* mediator that may contribute to CF airway inflammation.

## Introduction

Chronic infection by *Pseudomonas aeruginosa* is a major driver of adverse clinical outcomes in people with cystic fibrosis (pwCF) (1). Although highly effective modulator therapy reduces the relative abundance of *P. aeruginosa* and leads to substantial clinical improvements, persistent infections continue to occur in many pwCF, underscoring the need for continued vigilance (2, 3). Long-term colonization of the CF lung arises from both intrinsic host defects, such as viscous mucus, impaired mucociliary clearance and dysregulated immunity, and the remarkable ability of *P. aeruginosa* to adapt and evade immune clearance (4, 5). The extensive genomic variability, metabolic adaptability and phenotypic diversity of *P. aeruginosa* have contributed to its effectiveness as a dominant opportunistic pathogen in the CF airway (6-8).

During chronic infection, *P. aeruginosa* adapts to the CF lung by becoming less invasive and less virulent, enabling bacterial persistence while at the same time limiting extensive host damage (9). To this end, *P. aeruginosa* downregulates several virulence factors, including proteolytic activity, type III secretion system (T3SS) and motility, and switches to a sessile biofilm-based and mucoid lifestyle that enhances tolerance to both host defense systems and antibiotics (4, 9, 10). Additional adaptations include metabolic reprogramming to sustain growth under hypoxic mucus conditions and to use alternative nutrient sources, as well as selection of *P. aeruginosa* strains with mutations in quorum-sensing regulators such as *lasR*, and the emergence of hypermutator lineages and persister cells capable of long-term survival (4, 10).

A paradox in chronic pseudomonal infection in pwCF is that, despite attenuated virulence, exacerbated airway inflammation persists – a phenomenon that remains poorly understood. Studies have demonstrated that late-stage *P. aeruginosa* CF isolates trigger robust cytokine release *in vitro, in vivo* using animal models, and in pwCF (11-14). This excessive inflammatory response ultimately contributes to structural lung damage and a progressive decline in lung function (5, 15). While we recently highlighted the role of proteases (such as LasB) in immune evasion (16), the current study aims to identify previously unrecognized pro-inflammatory mediators involved in the excessive inflammation caused by *P. aeruginosa* during chronic infection in pwCF. To this end, *P. aeruginosa* was cultured under physiologically relevant conditions to stimulate the production of mediators likely expressed during *in vivo* infection. Specifically, we used synthetic CF sputum medium (SCFM2) to closely reproduce the nutritional environment of the CF lung and *P. aeruginosa* gene expression (17, 18). Furthermore, a three-dimensional (3D) alveolar epithelial cell model mimicking key phenotypic and functional characteristics of the native lung epithelium was used to assess host inflammatory responses (19-21). A comparative proteomics analysis of a previously generated dataset was performed to characterize differences between pro- and anti-inflammatory CF isolates, while existing *P. aeruginosa* CF sputum transcriptomic datasets were leveraged to confirm *in vivo* expression of the identified mediators.

In this study, we advance the understanding of mediators driving persistent inflammation during chronic *P. aeruginosa* CF lung infection and highlight potential targets for the development of novel therapeutic interventions.

## Materials & methods

### Bacterial species, culture conditions and supernatant preparation

*P. aeruginosa* isolates were previously obtained from the sputum samples of two individuals with CF (referred to as patient 1 and 6) (22). Patient 1 had an early-stage infection (1 year), whereas patient 6 had a long-term chronic infection (> 19 years). Throughout this paper, isolates are denoted as X/Y, where X is the isolate number and Y is the patient number. For example, 7/6 indicates isolate 7 from patient 6. In addition, several *P. aeruginosa* reference strains were included namely, PAO1, AA44 (a late CF sputum isolate), and AMT 0023-30, a pediatric early CF isolate (23, 24). AMT 0023-30 was used as a positive control for quantification of pyocyanin production as it has been reported as a high producer, while PAO1 was used as a positive control for the quantification of pyoverdine (23, 25). For select experiments, PA14 WT and PA14 Transposon (Tn) mutants derived from a *P. aeruginosa* transposon-mutant library were also used (24, 26, 27).

*P. aeruginosa* strains were cultured as described previously (16). In short, strains were grown in Luria-Bertani (LB) broth overnight at 37 °C with shaking at 250 revolutions per minute (rpm). All overnight *P. aeruginosa* cultures (PA14 WT, PA14 Tn mutants, PAO1, AMT 0023-30, AA44 and the 7 clinical isolates from patient 6) were diluted in SCFM2 to an OD_590_ of 0.005, equivalent to 5 × 10^7^CFU/mL, then further diluted to reach a bacterial cell density of 5 x 10^5^ CFU/mL. SCFM2 was prepared as described previously (17), except that mucin was autoclaved instead of UV-sterilized. Bacterial suspensions were incubated for 48 h under microaerophilic conditions (5.5–12 % O_2_) using Oxoid™ CampyGen™ Compact Sachet (CN0020C, Thermo Fisher Scientific) sealed with a plastic pouch to mimic oxygen-restricted conditions observed in pwCF with advanced lung disease (28). The medium control (MC) i.e., SCFM2 without bacteria, was incubated in parallel. Afterwards, the bacterial suspensions and the medium control were centrifuged (3500 rpm, 10 min) and the resulting supernatants were passed through a 0.22 µm filter to collect the cell-free supernatant. Supernatants were stored at – 20 °C without repeated freeze/thaw cycles.

Cell-free supernatants of PA14 Tn mutants were tested in the presence or absence of 0.5% (v/v) LasB inhibitor **4b**. This phosphonic acid inhibitor (C_13_H_16_F_3_NO_4_P, MW: 338.08 g/mol) was previously developed by *Konstantinović et al.* (29). The compound was first dissolved in DMSO, and the final concentration was adjusted to 0.5% (v/v) to minimize potential solvent effects of DMSO on bacterial culture supernatants, as previously described (29).

### Quantification of P. aeruginosa virulence factors

#### Pyoverdine measurement

Pyoverdine was quantified by transferring 200 μL of filtered, cell-free *P. aeruginosa* supernatant in duplicates into a flat-bottom 96-well plate. Absorbance was measured at 400 nm using the Victor® Nivo™ Multi-mode plate reader (Perkin Elmer, Shelton, CT, USA). PAO1 was used as a positive control.

#### Pyocyanin assay

Pyocyanin quantification was performed via the chloroform-HCl extraction method (30). Briefly, 1.5 mL of filtered cell-free *P. aeruginosa* supernatant was mixed with 1.5 mL chloroform by vortexing, followed by centrifugation (5000 rpm, 10 min) to separate the aqueous and organic phases. The chloroform layer was then collected and mixed with 0.2 N HCl. After another round of vortexing and centrifugation, 300 μL of the resulting pink aqueous (HCl) phase was transferred to a flat-bottom 96 well-plate, and absorbance was measured at 520 nm with the EnVision Multilabel Plate Reader (Perkin Elmer, Shelton, CT, USA). In parallel, the chloroform-HCl extraction method was applied to the supernatant of the medium control (i.e., SCFM2 alone) and AMT 0023-30, which served as the negative and positive controls, respectively.

#### Rhamnolipid quantification

Semi-quantitative detection of rhamnolipid production was performed using agar plates containing cetyltrimethylammonium bromide (CTAB) and methylene blue (MB) (31-33). Minimal medium agar plates were supplemented with 0.2 g/L CTAB and 5 mg/L MB. Holes were created into the plates and inoculated with 100 μL of a 5 x 10^7^ CFU/mL bacterial suspension in SCFM2. Plates were incubated at 37 °C under microaerophilic conditions (3% O_2_, 5% CO_2_ and 92% N_2_) using a hypoxia chamber (Bactrox, Sheldon manufacturing Inc., Cornelius, OR, US) for 48 h. Halo diameters surrounding the holes were measured to quantify rhamnolipid production. The medium control (100 μL) and 100 mM Tween-80 (100 μL) were used as the negative and positive controls, respectively.

#### Endotoxin quantification

Endotoxin levels in the filtered supernatants were measured using the Pierce Chromogenic Endotoxin Quant Kit (Thermo Fisher Scientific) following manufacturer’s instructions. Lipopolysaccharide (LPS) from *P. aeruginosa* serotype 10 (1 mg/mL; Merck, Darmstadt, Germany) was used as a positive control. Samples were diluted with Endotoxin-Free Water to achieve reading within the linear area of the standard curve (0.1–1.0 EU/mL), specifically a dilution factor of 10^5^ was used for bacterial cell-free supernatants and 5 x 10^6^ for the positive control. Each biological replicate was analyzed in two technical replicates.

#### Proteolytic and elastolytic activity assays

The proteolytic activity was assessed using the azocasein assay, while elastase activity was measured with the elastin-Congo red assay. Both assays were performed as described previously (16), with minor modifications. For each assay, 250 μL azocasein solution or elastin-Congo red suspension was combined with 250 μL of GTSF-2 medium containing 40 % (v/v) filtered cell-free bacterial supernatant, with and without the addition of 0.5% (v/v) component **4b** (final concentration 50 μM).

For the azocasein assay, samples were incubated at 37 °C for 1 h under shaking conditions (250 rpm). The reaction was stopped by adding 100 μL of 10% (w/v) trichloroacetic acid, followed by centrifugation at 13,000 rpm for 15 min. From the resulting supernatant, 100 μL was transferred to a flat-bottom 96-well plate and mixed with 100 μL 625 nM NaOH.

For the elastin-Congo red assay, samples were incubated at 37 °C for 24 h under shaking conditions (250 rpm), followed by centrifugation at 13000 rpm for 15 min. From the resulting supernatant, 200 μL was transferred to a flat-bottom 96-well plate. To avoid saturated absorbance reading, the supernatant was diluted in Milli-Q (MQ) water when necessary.

Finally, the OD was measured at 420 nm and 492 nm for the azocasein and elastin-Congo red assays, respectively, using the Victor® Nivo™ Multi-mode plate reader (Perkin Elmer). The same procedure was applied to the medium control mixed with either azocasein solution or Elastin-Congo red suspension, which served as the negative control. Two technical replicates were performed for each biological replicate.

### Arbitrary PCR and sequencing to confirm the identity of transposon mutant

Arbitrary polymerase chain reaction (PCR) and sequencing were performed as described previously (24, 26, 27). A culture derived from a purified *P. aeruginosa* PA14 Tn mutant colony was grown statically in 96-well plates containing 280 µL LB supplemented with 15 µg/mL gentamicin per well at 37 °C for approximately 40 h. Following incubation, 70 µL of culture was transferred to PCR tubes and stored at – 20 °C. For DNA extraction, samples were thawed, lysed at 99 °C for 10 min, and centrifuged to pellet cell debris (3,500 rpm, 5 min). For the first round of arbitrary PCR (ARB1), 3 µL of lysate was used as template, and amplification was performed with the transposon-specific primer PMFLGM.GB-3a and arbitrary primer ARB1D (***Table 1***), and the Q5® Hot Start High-Fidelity 2X Master Mix (New England Biolabs, Ipswich, MA USA) according to manufacturer’s instructions. Thermocycling conditions were as follows: 95 °C for 5 min; 30 cycles of 95 °C for 30 s, 47 °C for 45 s, and 72 °C for 1 min; followed by a final extension at 72 °C for 5 min. For the second round of arbitrary PCR (ARB2), 5 µL of ARB1 reaction was used as template. Reactions were carried out with the transposon-specific primer PMFLGM.GB-2a and arbitrary primer ARB2A (***Table 1***), and the Q5® Hot Start High-Fidelity 2X Master Mix under the following conditions: 40 cycles of 95 °C for 30 s, 45 °C for 30 s and 72 °C for 1 min; followed by a final extension at 72 °C for 5 min. PCR products were purified by mixing 5 μL ARB2 reaction with 2 μL ExoSAP-IT reagent (Thermo Fisher Scientific) and processed according to the manufacturer’s protocol. Sanger sequencing was performed after mixing the purified products (10 ng/μL) with the sequencing primer (5 μM) (***Table 1***; LightRun Tube Service; Eurofins Genomics). The resulting sequences were then queried against the *P. aeruginosa* UCBPP-PA14 genome (taxid:208963) using the BLASTx algorithm implemented through the NCBI Basic Local Alignment Search Tool (34-36).

**Table 1:**
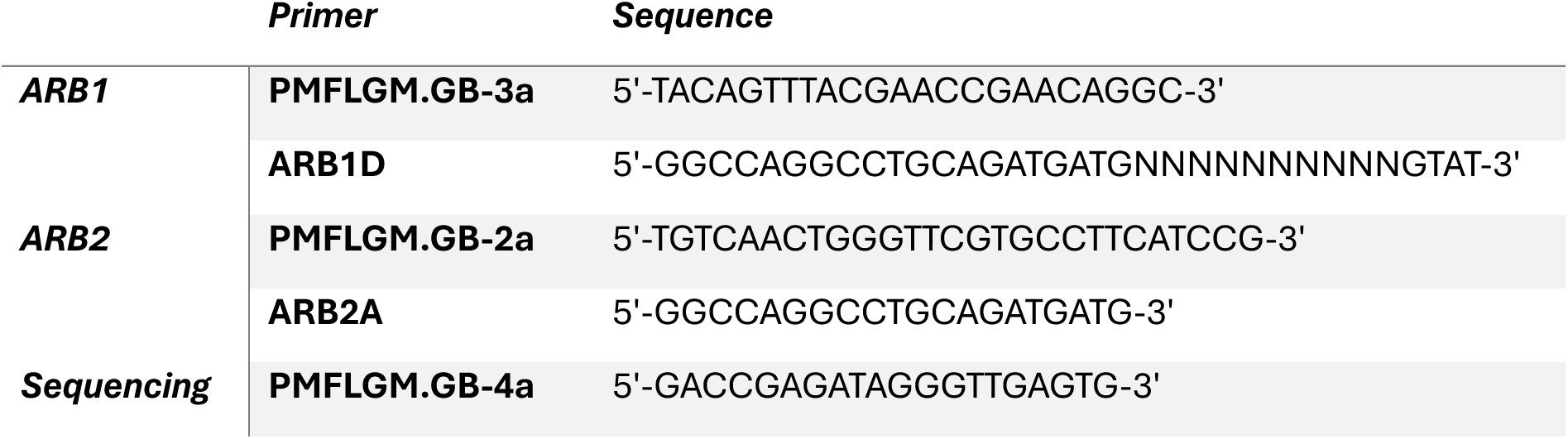
Sequences of used primer pairs.

### Three-dimensional lung epithelial cell culture model

The human adenocarcinomic alveolar epithelial cell line A549 (ATCC, CCL185) was cultured as an organotypic 3D cell culture model using a rotating wall vessel (RWV) bioreactor, as described previously (20, 21). Briefly, A549 cells were maintained as monolayers in T75 flasks using GTSF-2 medium (Hyclone™, Logan, UT, USA) supplemented with 1.5 g/L sodium bicarbonate (Sigma-Aldrich), 10% heat-inactivated fetal bovine serum (FBS) (Thermo Fisher Scientific), 2.5 mg/L insulin-transferrin-sodium selenite (Sigma-Aldrich) and 1% penicillin-streptomycin (with a stock concentration of 10,000 units/mL penicillin and 10 mg/mL streptomycin; Sigma-Aldrich). Upon reaching confluence, cells were washed with Hank’s Balanced Salt Solution (HBSS) (Life Technologies, Carlsbad, CA, USA), and dissociated using 0.25% trypsin-EDTA (Thermo Fisher Scientific). Cell counts and viability were assessed via trypan blue (0.4%Sigma Aldrich) using a hemocytometer. A suspension containing 2 x 10^6^ viable cells in supplemented GTSF-2 medium was combined with 0.25 g porcine-skin collagen-coated dextran beads (Cytodex®-3 microcarrier beads, Cytiva, Marlborough, MA, USA), transferred to RWV bioreactors – first in a slow turning lateral vessel (STLV) and after 1 week in a high-aspect rotating vessel (HARV) (Synthecon, Houston, TX, USA) – and maintained for 11–14 days to allow the formation of differentiated 3D lung epithelial aggregates. All cultures were incubated at 37 °C with 5% CO_2_.

### In vitro cell-exposure assay

3D A549 aggregates were transferred to a flat-bottom 96-well plate at a density of 2.5 x 10^5^ cells per well in 150 µL GTSF-2 medium without FBS and antibiotics. Host cells were exposed to 100 µL of either cell-free supernatant from *P. aeruginosa* cultures or filtered medium control (SCFM2) for 4 h at 37 °C under microaerophilic conditions (3% O_2_, 5% CO_2_ and 92% N_2_) using a hypoxia chamber (Bactrox, Sheldon manufacturing Inc.). The exposure experiment using cell-free supernatant of PA14 Tn mutant was performed in the absence or presence of 0.5% (v/v) component **4b** (final concentration 50 μM). As a positive pro-inflammatory control, 25 μL of supernatant of *P. aeruginosa* strain AA44 was used. Following incubation, the conditioned medium (i.e., the supernatant fraction from the 3D lung cells after exposure to bacterial cell-free supernatant) was collected and stored at – 20 °C until further analyses, avoiding repeated freeze/thaw cycles.

### Cytotoxicity assay

Previous work demonstrated that *P. aeruginosa* protease production interferes with conventional lactate dehydrogenase (LDH) quantification (37). To overcome this, a modified protocol was applied (37). In this approach, LDH release from the viable cell fraction remaining adherent to the microcarrier beads at the end of the exposure experiment is measured as an indicator of cell viability. Briefly, cultures were rinsed twice with HBSS, and adherent cells were lysed with 1% Triton X-100 (Sigma-Aldrich). LDH activity was subsequently quantified using the Lactate Dehydrogenase Activity Assay Kit (Sigma-Aldrich) according to the manufacturer’s instructions.

### Inflammatory marker quantification

For *in vitro* cell exposure assays, Interleukin (IL)-8 release was measured in the conditioned medium using the Human ELISA MAX^TM^ Standard Set (Biolegend) according to the manufacturer’s instructions. Samples were diluted in GTSF-2 medium without FBS and antibiotics.

### Proteomics analyses of the culture supernatants of clinical P. aeruginosa isolates grown in SCFM2

#### Comparative proteomics analysis & data visualization

Untargeted proteomics of the cell-free supernatant from all isolates included in this study was previously performed in (16) and deposited in the ProteomeXchange Consortium via the PRIDE partner repository with the dataset identifier PXD065321. A new comparative proteomics data analysis was performed in the present study (comparing different isolate groups as previously), hereby comparing 5 pro- with 6 anti-inflammatory *P. aeruginosa* CF isolates (listed in ***Table 2***). Methodological details for sample preparation, liquid chromatography-tandem mass spectrometry (LC-MS/MS) acquisition, and spectral data analysis are available in (16). Isolates were categorized into pro-inflammatory and anti-inflammatory groups based on their significant induction/reduction of IL-8 secretion by 3D lung epithelial cells (***Table 2 & Figure S.1***).

**Table 2:** Sample groups according to isolate and inflammatory profile.

| Sample name | Isolate | Inflammatory response | Number of samples |
| --- | --- | --- | --- |
| 6/1 rep1-rep2-rep3-rep4 | 6/1 | Anti-inflammatory | 4 |
| 12/1 rep1-rep2-rep3 | 12/1 | Anti-inflammatory | 3 |
| 14/1 rep2-rep3-rep4 | 14/1 | Anti-inflammatory | 3 |
| 15/1 rep2-rep3-rep4-rep1 | 15/1 | Anti-inflammatory | 4 |
| 16/1 rep2-rep3-rep4-rep1 | 16/1 | Anti-inflammatory | 4 |
| 19/1 rep2-rep3-rep4 | 19/1 | Anti-inflammatory | 3 |
| 6/6 rep1-rep2-rep3-rep4 | 6/6 | Pro-inflammatory | 4 |
| 9/6 rep4-rep1-rep3 | 9/6 | Pro-inflammatory | 3 |
| 12/6 rep4-rep1-rep2 | 12/6 | Pro-inflammatory | 3 |
| 13/6 rep3-rep4-rep1 | 13/6 | Pro-inflammatory | 3 |
| 14/6 rep1-rep2-rep3 | 14/6 | Pro-inflammatory | 3 |

For differential expression analysis, missing values were imputed from a normal distribution using default parameters in Perseus (38). Group comparisons were performed using a two-sample t-test with permutation-based false discovery rate (FDR) set at 5%.

#### Functional annotation

The gene names corresponding to the 605 significantly upregulated proteins (*p* < 0.05, fold change > 2) in the pro-inflammatory isolate group were uploaded to the Database for Annotation, Visualization, and Integrated Discovery (DAVID) (search performed on December 5, 2024) (39, 40). Functional annotation was performed using *P. aeruginosa* PAO1 as the reference strain. Based on annotation terms linked to established virulence processes and/or PubMed text mining of the respective factors, 62 proteins were identified as being associated with virulence (***Table S.1***).

#### Prediction of protein subcellular localization

The subcellular localization of mediators of interest was retrieved from the Pseudomonas Genome Database (version 22.1) with *P. aeruginosa* PAO1 (Stover *et al.,* 2000) as reference strain (41, 42).

### Analysis of RNA-seq datasets of P. aeruginosa transcriptomes in CF sputum

Transcriptomic data for *P. aeruginosa*, including variance-stabilized (VST) normalized count data from sputum samples of pwCF were obtained from the supplementary materials of Lewin *et al*. (43). To identify additional publicly available transcriptomic datasets, we accessed a comprehensive pre-release *P. aeruginosa* database (v0.1.0-beta) maintained by the Whiteley Lab at Georgia Institute for Technology (https://www.thewhiteleylab.com/database). This resource provides normalized gene expression data expressed as transcripts per million (TPM), along with the corresponding sample metadata. The downloaded input for the application included feature counts, data summaries, and metadata (dated: July 18, 2025). Filtered results of the transcriptomic dataset (from the original study Rossi *et al.* (44); SRA: ERP106536) were saved as CSV files for subsequent analyses.

### Statistical analysis

ELISA data were processed using GainData® (Arigo Biolaboratories, available at https://www.arigobio.com/elisa-analysis). Standard curves were fitted using a four-parameter logistic regression model, and cytokine concentrations were calculated from absorbance values within the linear region of the curve.

Statistical analyses of virulence factor quantification, LDH cytotoxicity assay, and cytokine concentrations of 3D aggregates exposed to PA14 WT and PA14 Tn mutants were performed in GraphPad Prism (Version 10; https://www.graphpad.com). Data distribution was assessed using the Shapiro–Wilk test. Non-normally distributed data were analyzed using Mann–Whitney U or Kruskal–Wallis tests. To account for multiple comparisons, FDR correction was applied using the Benjamini–Hochberg procedure, with a significance set at 5% (45). Paired data for proteolytic and elastolytic activities of the PA14 Tn mutants, measured in the absence or presence of compound **4b**, were analyzed using a mixed-effects model with restricted maximum likelihood (REML) estimation.

All experiments were performed in at least three biological replicates. Figure and data visualizations were generated using GraphPad Prism and R (Version 4.4.2), employing R packages such as ggplot2 (46, 47).

## Results

### Identification of candidate pro-inflammatory mediators of P. aeruginosa using comparative proteomics analysis

In our previous study, we observed that the supernatant obtained from *P. aeruginosa* CF isolates cultured in SCFM2 induced a variable inflammatory response based on IL-8 secretion in an organotypic 3D lung cell culture model. In particular, some isolates showed a pro-inflammatory response while others exhibited an anti-inflammatory response (***Figure S.1)***. All isolates causing robust pro-inflammatory effects in the 3D lung model were obtained from a single sputum sample from an individual with CF with long-term (over 19 years) chronic *P. aeruginosa* infection, while isolates leading to an anti-inflammatory response were derived from an individual with CF who had become chronically infected relatively recently (1 year). Our previous work showed that proteolytic and elastolytic activity (partially driven by the metalloprotease elastase B, LasB) mediate the anti-inflammatory effect of CF isolates through cytokine degradation (16). In the present study, we aimed to identify pro-inflammatory mediators produced by CF isolates with pro-inflammatory activity.

To this end, a dual strategy was employed: 1) investigating the potential involvement of well-known virulence factors – pyocyanin, pyoverdine, rhamnolipids, and endotoxins – using semi-quantitative assays; 2) performing a comparative proteomics analysis of a previously generated dataset to characterize differences between pro-inflammatory and anti-inflammatory isolate groups (16).

Firstly, for all pro-inflammatory CF isolates tested, no significant production of any of the four well-established virulence factors was detected (***Figure S.2.A, S.2.B, S.2.C and S.2.D***). Some CF isolates showed endotoxin levels of up to approximately 20,000 EU/mL, which, nevertheless, was not statistically significantly different from the medium control (SCFM2 alone) (***Figure S.2.D***). Furthermore, no significant correlation was observed between the endotoxin concentration in the cell-free supernatants and IL-8 release (Spearman’s correlation coefficient r = 0.0476; p = 0.9349; ***Figure S.2.E***). Hence, well-known virulence factors of *P. aeruginosa* are likely not responsible for the observed pro-inflammatory effect.

Secondly, to identify previously unrecognized pro-inflammatory mediators of *P. aeruginosa*, we performed a comparative analysis of shotgun proteomics data from supernatants of CF isolates belonging to each of the inflammatory groups, after culturing in SCFM2 (16). Data analysis was performed on eleven clinical isolates, classified into 6 anti-inflammatory isolates and 5 pro-inflammatory isolates (**Table 2**). In total, 2,196 protein groups were quantified. A two-sample t-test between the pro-inflammatory and anti-inflammatory isolate group was performed using a permutation-based FDR of 5% to correct for multiple testing. The results are presented in the volcano plot shown in ***Figure 1***. Of the 1,181 significantly differentially expressed proteins (*p* < 0.05, fold change > |2|), a total of 629 were upregulated in the pro-inflammatory isolates group, while 552 proteins were downregulated. The gene names corresponding to the 629 upregulated proteins in the pro-inflammatory isolate group were uploaded to DAVID to perform functional annotation. Based on annotation terms linked to established virulence processes, 62 proteins were identified as being associated with virulence (***Table S.1***). Additionally, we observed a downregulation of multiple extracellular proteases – including LasB, alkaline protease A (AprA), and protease IV (PrpL) – alongside several proteins associated with Type IV pilus (PilV, PilW, PilY1, PilY2, PilE) and flagellum-dependent motility (FliC, FlgK, EstA, FlgE, CheZ) (***Figure S.3***).

**Figure 1:**
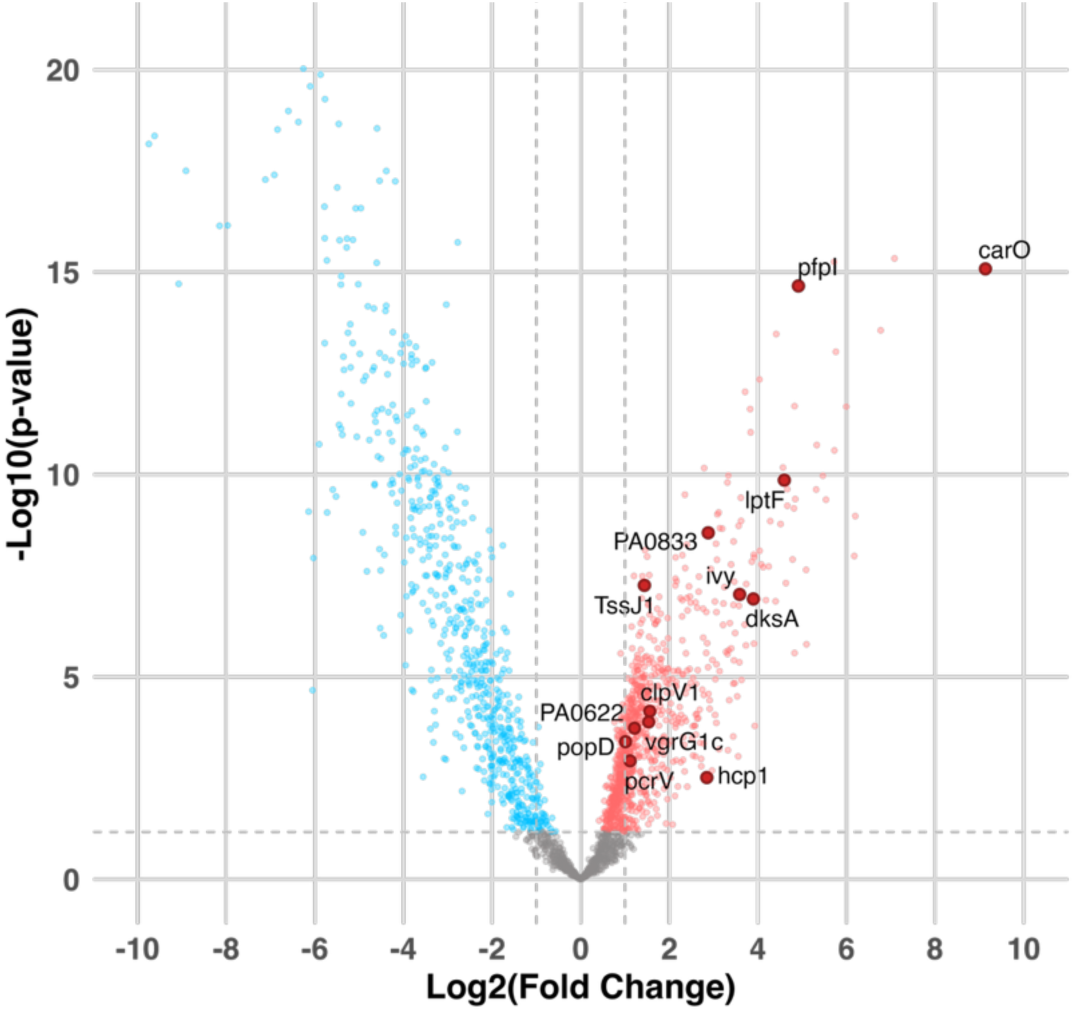
Volcano plot with differentially expressed proteins of interest in the pro-inflammatory isolate group: A volcano plot was generated to display the differences in protein abundance between the pro- and anti-inflammatory isolate group. The difference is represented as log_2_(Fold Change), plotted against -log_10_(p-values) derived from a two-sample t-test. Symbols corresponding to proteins of interest are enlarged and labeled. The horizontal dashed line indicates the significance threshold, determined using a permutation-based FDR correction of 5%.

As a next step, we selected candidate pro-inflammatory mediators from the proteins upregulated in the pro-inflammatory isolate group for downstream *in vitro* validation of inflammatory activity. Candidates were selected based on functional annotation or PubMed text-mining evidence linking them to virulence, and/or strong upregulation (fold change > 5), as well as the availability of corresponding *P. aeruginosa* PA14 mutants in a Tn mutant library. Notably, the genes corresponding to some of the candidate pro-inflammatory mediators were not inactivated in the Tn mutant library, or did not pass the arbitrary PCR quality control. In total, 13 mediators meeting these criteria were selected for further analysis (***Figure 1 & Table 3***).

**Table 3:** Selection of candidate pro-inflammatory proteins. Subcellular localization is available in the Pseudomonas Genome Database (https://www.pseudomonas.com/). Abbreviations: Outer Membrane Vesicle (OMV), Type III secretion system (T3SS), Type IV secretion system (T6SS)

| Gene | PA Locus | Gene Name | Function and their association with virulence and/or inflammation (if available) | Localization | Ref. |
| --- | --- | --- | --- | --- | --- |
| **carO** | PA0320 | Calcium-regulated OB-fold protein CarO | Maintaining intracellular Ca<sup>2+</sup> homeostasis | Unknown | (48) |
| **dksA** | PA4723 | Suppressor protein DksA | Transcriptional regulator, stringent response modulator | Cytoplasmic | (49-53) |
| **ivy** | PA3902 | Hypothetical protein | Inhibitor of lytic transglycosylase activity and vertebrate lysozyme | Periplasmic | (54, 55) |
| **vgrG1c** | PA2685 | Type VI secretion system spike protein VgrG1c | Secretion machinery protein of the contractile tail tube complex from the T6SS | Cytoplasmic | (56) |
| **pcrV** | PA1706 | Type III secretion protein PcrV | Needle tip protein of the syringe-like injectisome T3SS | Extracellular | (57, 58) |
| **TssJ1** | PA0080 | Hypothetical protein | The T6SS consists of a membrane-anchored apparatus containing the outer membrane lipoprotein TssJ | Cytoplasmic<br>Membrane<br>OMV | (59) |
| **clpV1** | PA0090 | Secretion protein ClpV1 | an AAA + ATPase, which supplies energy to the T6SS. Its loss completely abolishes secretion activity | Cytoplasmic | (60) |
| **hcp1** | PA0085 | Protein secretion apparatus assembly protein | Secretion machinery protein of the contractile tail tube complex from the T6SS | Extracellular | (56, 57) |
| **popD** | PA1709 | Translocator outer membrane protein PopD | PopD contributes to the formation of a translocation pore in the host cell membrane by the T3SS | Extracellular | (61) |
| **lptF** | PA3692 | Lipotoxin F | Outer membrane protein that is highly expressed in mucoid *P. aeruginosa* isolates from chronic CF infections, survival factor, stimulates inflammatory responses, promising vaccine candidate | Outer Membrane<br>OMV | (62-64) |
| **PA0833** | PA0833 | OmpA-like domain-containing protein | Outer membrane protein, promising vaccine candidate | Outer Membrane<br>OMV | (65, 66) |
| **pfpl** | PA0355 | Protease Pfpl | Intracellular protease associated with antibiotic susceptibility, swarming motility, and biofilm formation | Cytoplasmic | (67) |
| **PA0622** | PA0622 | Probable bacteriophage protein | A protein of unknown function that is highly abundant in OMVs from *P. aeruginosa* biofilms | Unknown<br>OMV | (68-70) |

### In vitro validation of candidate pro-inflammatory mediators

To evaluate the role of the selected mediators in modulating host inflammatory responses *in vitro*, IL-8 release was measured in an organotypic 3D lung cell culture model following exposure to cell-free supernatants from PA14 Tn mutant strains deficient in each mediator.

The *P. aeruginosa* PA14 WT and Tn mutants are strong LasB producers (***Figure S.4.A & S.4.B***). A strong downregulation of LasB was observed in the pro-inflammatory isolates group, consistent with previous observations (***Figure S.3***) (16). Therefore, we inhibited LasB activity in the cell-free supernatants of the PA14 Tn mutant strains by addition of the LasB-specific phosphonic acid derivative **4b** (29), and subsequently quantified IL-8 release by the 3D lung aggregates.

Initially, the activity of compound **4b** was validated in our experimental setup using a dose-response study. Mixed-effects modeling with REML estimation revealed a significant inhibitory effect of compound **4b** (50 μM) on both proteolytic and elastolytic activities of the selected PA14 transposon mutants (p < 0.001 for each activity; ***Figure S.4.A & S.4.B***). Treatment with the inhibitor resulted in a significant increase in IL-8 release by 3D lung aggregates exposed to PA14 Tn mutant cell-free supernatants, indicating restored cytokine activity due to reduced LasB-mediated degradation (***Figure S.5***). Comparative analysis of IL-8 release induced by each Tn mutant strain and the PA14 WT demonstrated a significant anti-inflammatory effect only for PA14 dksA::Tn, but not for other mutants (***Figure 2***).

**Figure 2:**
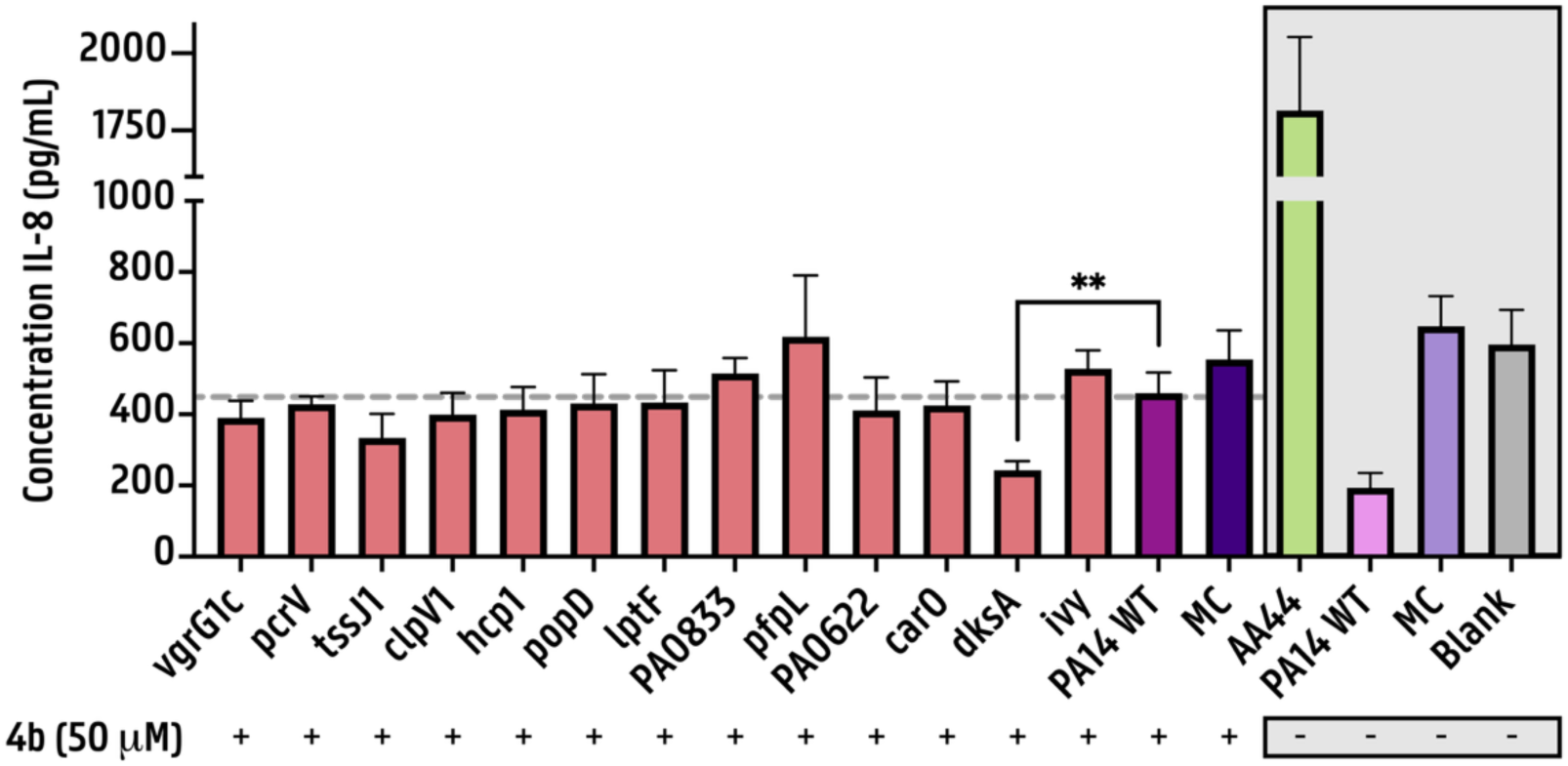
Inflammatory response triggered by cell-free supernatants of PA14 Tn mutants, evaluated in the presence of LasB inhibition: IL-8 release from an organotypic 3D lung cell culture model following 4 h exposure to 40% (v/v) P. aeruginosa cell-free supernatants, in the presence of 50 μM LasB inhibitor **4b** (0.5% (v/v)). SCFM2 alone was used as an MC. The blank represents untreated A549 cells cultured in GTSF-2 medium without FBS. The blank and MC represent the basal level of cytokines that the cells produce in the absence of a pro-inflammatory stimulus. (n = 3–6, statistical analysis was performed using the non-parametric Kruskal–Wallis ANOVA comparing to the PA14 WT with inhibitor, followed by the Benjamini–Hochberg correction for multiple testing with an FDR of 5%, * p < 0.05, ** p < 0.01, *** p < 0.001, error bars represent the standard error).

Cell-viability assays were conducted to determine whether potential cytotoxic effects contributed to the observed results. Neither the bacterial supernatants nor the compound alone or in combination caused significant cytotoxicity in any experimental condition (***Figure S.6.A & S.6.B***).

### DksA-dependent regulation of virulence factors and the proteome

To investigate the reduced inflammatory activity of PA14 dksA::Tn, we assessed key pro-inflammatory virulence factors, specifically rhamnolipid production and proteolytic activity, as decreases in both have been linked to *dksA* mutations (49). While no statistically significant differences were observed for the tested virulence factors, a clear reduction in rhamnolipid production was observed in PA14 dksA::Tn as compared to the wild-type strain (***Figure S.7.A, S.7.B & S.7.C)***.

Given that DksA is a global transcriptional regulator influencing over 1,500 genes (52), we next examined the subset of DksA-regulated proteins that were differentially expressed exhibiting a fold change greater than |2| (as identified by our comparative proteomics analysis). This approach revealed 385 DksA-regulated proteins in our dataset (***Figure S.8 and Table S.2***), including several of the candidate pro-inflammatory proteins: CarO, PcrV, PopD, LptF and PfpI (***Table 3***). Additionally, we identified multiple virulence-associated proteins, such as components of flagellar assembly (FlgE, FlgK, FliC), and type IV pilus biogenesis (pilV, pilW, pilY1, pilY2, and pilE), highlighting the wide impact of DksA on pathogenicity-related processes in *P. aeruginosa*.

### Validating the expression of dksA and other selected mediators in human CF sputum

Next, we evaluated the relevance of our findings for the CF population, by assessing the expression of *P. aeruginosa* pro-inflammatory genes in sputum of pwCF. Initially, we leveraged a transcriptomic dataset consisting of 24 *P. aeruginosa* sputum transcriptomes collected from 21 pwCF at two CF clinics, one in Copenhagen, Denmark, and the other in Atlanta, Georgia, USA, previously reported by Lewin *et al.* (43). Genes were ranked by expression across CF sputum samples to identify those most highly expressed (***Figure 3.A and 3.B***). Notably, from our 13 selected mediators, *dksA*, *vgrG1c*, and *PA0833* ranked among the top 25% expressed genes. Normalized count data for *PA0622* was absent from the dataset as Lewin *et al.* excluded this gene due to its expression in only 14 of the 24 sputum samples. With the exception of *carO*, (detected in 90% of samples), *pcrV* and *popD* (each detected in 95%), all other mediators were expressed in 100% of the sputum transcriptomes (43).

**Figure 3:**
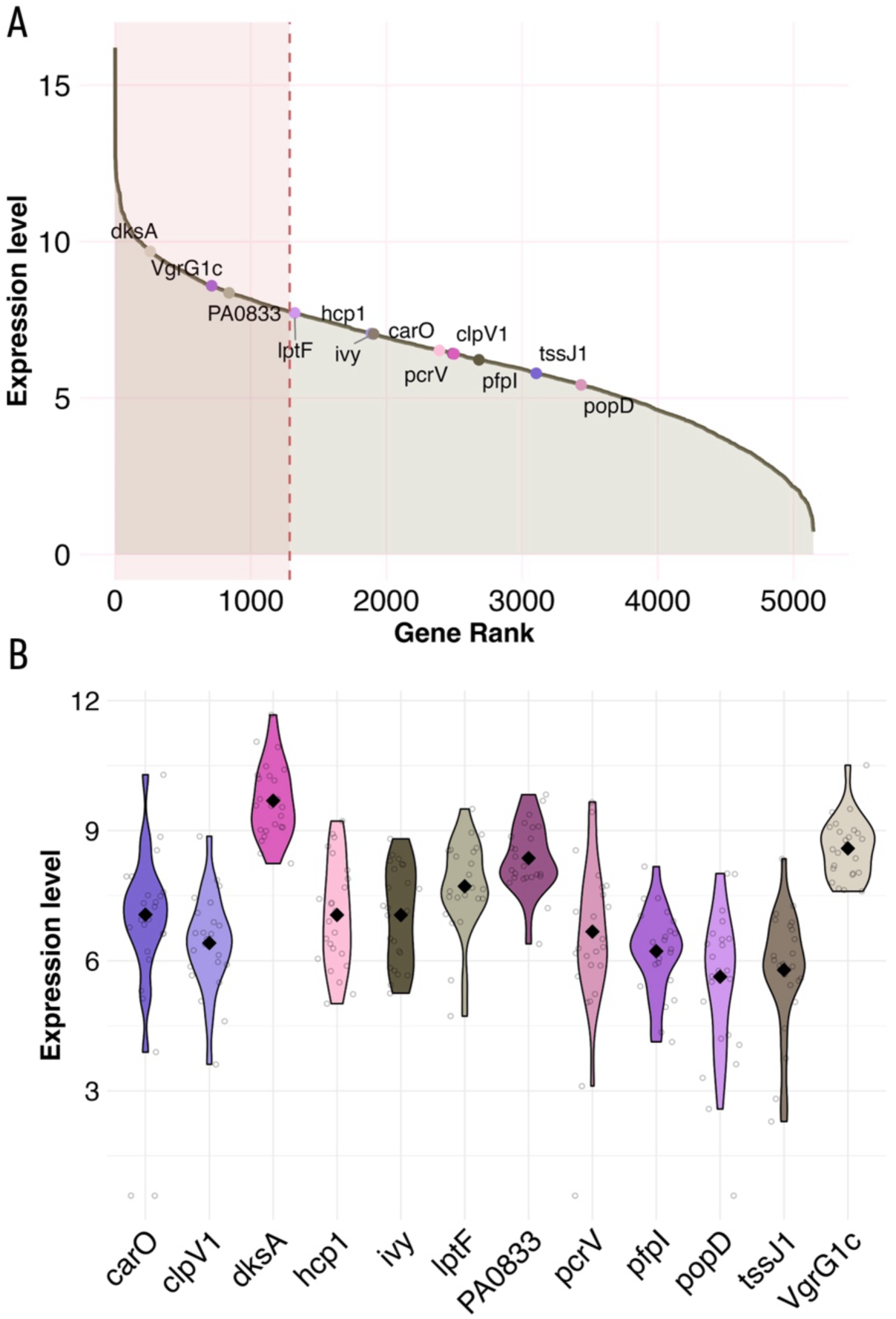
Expression profile of mediators of interest across 24 P.aeruginosa transcriptomes derived from CF sputum samples (data from Lewin et al. (43)): (A) Cumulative gene expression plot displaying genes ranked by their average expression (normalized count data) across the 24 CF sputum-derived P. aeruginosa transcriptomes. Proteins of interest are labeled, and the top quartile (25%) of most highly expressed genes is highlighted with a transparent red box; (B) Violin plots depicting the distribution and variability of expression levels for selected mediators across all 24 samples with each violin representing one gene. Samples with absent expression (zero counts) were represented as dots at baseline and excluded from the violin plot visualization.

In order to further investigate *P. aeruginosa* gene expression of the mediators in pwCF, we leveraged additional transcriptomic data using the *P. aeruginosa* database (https://www.thewhiteleylab.com/database). Data were leveraged from nine *P. aeruginosa* transcriptomes derived from CF sputum samples belonging to four different pwCF at different timepoints (originally derived from Rossi *et al*. (44)). These samples were collected from Copenhagen, Denmark, and all four pwCF were chronically infected with *P. aeruginosa* DK01 and/or DK02 lineage for more than 30 years. The expression profile of mediators of interest across all samples were visualized (***Figure S.10.A and S.10.B)*** and *dksA*, *VgrG1c*, *PA0833*, *carO* and *lptF* ranked among the top 25% expressed genes.

Given the confirmed role of DksA in the pro-inflammatory activity of *P. aeruginosa* (***Figure 2***), we further explored *dksA* expression in both transcriptomic datasets derived from CF sputum samples. Since *P. aeruginosa* encodes two functional *dksA* paralogs, we extended our analysis by including *dksA2*. Both *dksA1* (synonym: *dksA*) and *dksA2* regulators were among the top 25% of expressed genes in all sputum samples (***Figure S.9.A, S.9.B, S.10.C and S.10.D***; data from Lewin *et al.* (43) and Rossi *et al.* (44), respectively).

## Discussion

Despite the progressive loss of well-known virulence factors, late-stage CF isolates of *P. aeruginosa* have been shown to elicit strong pro-inflammatory cytokine responses *in vitro*, *in vivo* using animal models and in the lungs of pwCF (11-13). In this study, we aimed to identify previously unrecognized bacterial mediators sustaining this persistent inflammation by analyzing multiple pro-inflammatory isolates from an individual with CF with over 19 years of chronic *P. aeruginosa* infection.

Major virulence factors including pyocyanin, pyoverdine, rhamnolipids and LPS, were negligible in the secretome of pro-inflammatory isolates, excluding their role in the observed phenotype. Comparative proteomics analysis guided the selection of 13 upregulated mediators from these isolates for further investigation. Using *P. aeruginosa* PA14 Tn mutants for validation, we found that the transcriptional regulator *dksA* was involved in the pro-inflammatory response of this pathogen.

Indeed, DksA was a notable finding in the proteome of the cell-free supernatants, given its known intracellular localization. Its extracellular presence is most likely due to cell lysis, although incorporation into membrane vesicles (OMVs) or release during cytoplasmic leakage associated with OMV formation is also possible. Additionally, several cytosolic proteins are known to perform additional extracellular functions, such as promoting biofilm formation or enhancing virulence (68, 71). However, it remains unclear whether DksA possesses any distinct extracellular role or is secreted via OMVs, warranting further investigation.

Regardless, DksA post-transcriptionally regulates hundreds of genes, including quorum-sensing-dependent virulence genes (e.g., encoding rhamnolipids and elastase), genes involved in tolerance to H_2_O_2_-induced oxidative stress and protection against macrophage-mediated killing (52, 53). We confirmed in the PA14 dksA::Tn mutant that rhamnolipid production was controlled by DksA. Since the pro-inflammatory CF isolates tested in the present study showed no detectable rhamnolipid production (which is often observed for chronic isolates (72)), the observed immunomodulation by DksA is likely not mediated by these molecules.

A recent study by Weimann *et al*. (73) demonstrated that intracellular survival within CF macrophages is facilitated by DksA1. Moreover, the authors reported that both the expression of the stringent response modulator *dksA1* and the activation of its associated regulon were linked to CF-specific adaptation of *P. aeruginosa* (73).

Interestingly, *P. aeruginosa* encodes two functional *dksA* paralogs, *dksA1* and *dksA2*, which are largely interchangeable but exhibit optimal activity under different environmental conditions (52). The zinc-finger motif of zinc-dependent dksA1 becomes structurally unstable during zinc-depletion, whereas *dksA2* is exclusively expressed under zinc starvation (52, 74, 75). Transcriptomic analyses of *P. aeruginosa* in CF sputum revealed high expression of both paralogs in our study. However, considering zinc starvation conditions in the CF lung environment, the biological relevance of increased *dksA1* expression under these conditions requires further clarification (43, 44, 76). *Z*inc deprivation is a well-recognized host-defense mechanism, exacerbated in CF sputum by high levels of calprotectin, a neutrophil-derived zinc-chelating protein (76). When comparing *in vivo* transcriptomes of 12 CF sputum samples with *in vitro* stationary-phase LB cultures of *P. aeruginosa* PA14 and matched clinical isolates, Rossi *et al.* (44) observed strong induction of *dksA2* and repression of *dksA*1. These findings indicate that, during CF lung infection, zinc-independent *dksA2* predominates in mediating stress responses (44). In the present study, proteomics analysis did not reveal DksA2 in the culture supernatant of the clinical CF isolates, likely because the isolates were cultured in SCFM2, which is not zinc-limited. Indeed, when zinc limitation is established in SCFM2 by chelation with calprotectin, *dksA2* becomes expressed (43). Hence, while DksA1 was likely involved in the pro-inflammatory response of *P. aeruginosa* in the model systems used in our study, further investigation is needed to define the specific roles of DksA1 and DksA2 and their downstream gene regulatory networks in the *P. aeruginosa*-induced inflammation in the CF lung environment.

Furthermore, twelve other candidate mediators were explored for their role in the pro-inflammatory response, but the cell-free supernatant of each individual mediator PA14 Tn mutant did not significantly influence IL-8 release relative to the PA14 wild-type. Six mutants were associated with the T3SS or T6SS, which are virulence factors that operate via a one-step mechanism that injects toxic effector proteins into target cells (77, 78). Both secretion systems have been shown to trigger host inflammation by inducing cytokine production such as IL-6, IL-8, and IL-1β in A549 cells and macrophages (79-82). Furthermore, immunization with a trivalent vaccine containing PcrV (T3SS), OprI, and Hcp1 (T6SS) proteins conferred protection in a murine *P. aeruginosa* pneumonia model (57), and a phase I/II trial in pwCF demonstrated that the anti-PcrV antibody KB001-A was effective in reducing sputum IL-8 levels (58). However, in this study, none of the T3SS- or T6SS-related mutants induced altered inflammatory responses relative to the PA14 wild-type. A key difference between our study and previous work is that earlier studies infected cells with live bacteria, whereas we employed cell-free supernatants. Since the *P. aeruginosa* secretion systems are anchored to the outer membrane and require this association for activity, their activity is typically absent in cell-free supernatants. Nevertheless, *P. aeruginosa* secretes OMVs, which have been implicated in virulence-associated interactions with lung epithelial cells (83, 84). While the presence of T3SS or T6SS proteins in *P. aeruginosa* OMVs has not been definitively reported, T3SS effectors and translocon proteins have been identified in OMVs from other bacteria including *Salmonella enterica* and *Escherichia coli* O157:H7 (85-87).

Analyses of publicly accessible transcriptomic data of CF sputum revealed high expression of both *PA0833* and *lptF*. Notably, LptF (Lipotoxin F) is highly expressed in mucoid *P. aeruginosa* isolates from chronic CF infections (62), and along with PA0833, is secreted via OMVs (41, 42, 70, 88). Both proteins have emerged as promising vaccine candidates. Immunization with PA0833 has been shown to protect against *P. aeruginosa* in a pneumonia mouse model (65, 66), while LptF activated NF-κB in human respiratory epithelial cells via Toll-like receptor 2 (63). Additionally, recent immunoinformatic approaches have designed an epitope-based peptide vaccine based on LptF that shows high stability and immunogenicity *in silico* (64). Elucidating the specific contributions of both mediators to inflammation remains an interesting direction for future investigation.

This study reveals considerable variability in the inflammatory profile of *P. aeruginosa* isolates from pwCF, with some retaining strong inflammatory activity despite the loss of well-known virulence traits. Culturing these isolates under physiologically relevant conditions uncovered mediators that could play a role in the persistent inflammation in the lungs of pwCF. Interestingly, these mediators were consistently and robustly expressed across all analyzed *P. aeruginosa* CF sputum transcriptomes. While DksA was the only mediator for which a pro-inflammatory role could be confirmed, it is likely that it may act in an additive or synergistic way with other identified mediators. Hence, the pro-inflammatory response is probably driven by multiple mediators in concert rather than a single factor. Future studies should elucidate potential interactions among mediators underlying the inflammatory response, and identify those with the greatest promise as therapeutic targets to mitigate chronic inflammation in *P. aeruginosa* CF lung infections.

## Author contributions

AC, XS and MW conceptualized the study and designed the experimental setup. SVDB provided the clinical isolates originating from sputum samples from pwCF. MW and LB performed the experiments and data analysis. AC, FI, LG, EVB, MWh and AKHH provided guidance throughout the study. DVH, SDF, SDV and MW contributed to the comparative proteomics data analysis. MW and AC wrote the manuscript, with input from all authors. All authors contributed to the article and approved the submitted version.

## Data availability statement

No new data were generated in this study. The mass spectrometry-based proteomics data analysed here have been deposited in the ProteomeXchange Consortium via the PRIDE partner repository under the dataset identifier PXD065321. *P. aeruginosa* CF sputum transcriptome normalized count data from Lewin *et al*. (43) are available as Dataset S3 in the original publication. *P. aeruginosa* CF sputum transcriptome normalized data from Rossi *et al*. (44) were obtained from the *P. aeruginosa* database (v0.1.0-beta; https://www.thewhiteleylab.com/database).

## Acknowledgments

We thank Prof. Dr. Piet Cools for generously providing the *P. aeruginosa* PA14 wild-type and PA14 transposon mutant strains.

## Supplementary Data

The data presented in **Figure S.1** was reproduced from our previous work; no new experiments were conducted (16).

**Figure S.1:**
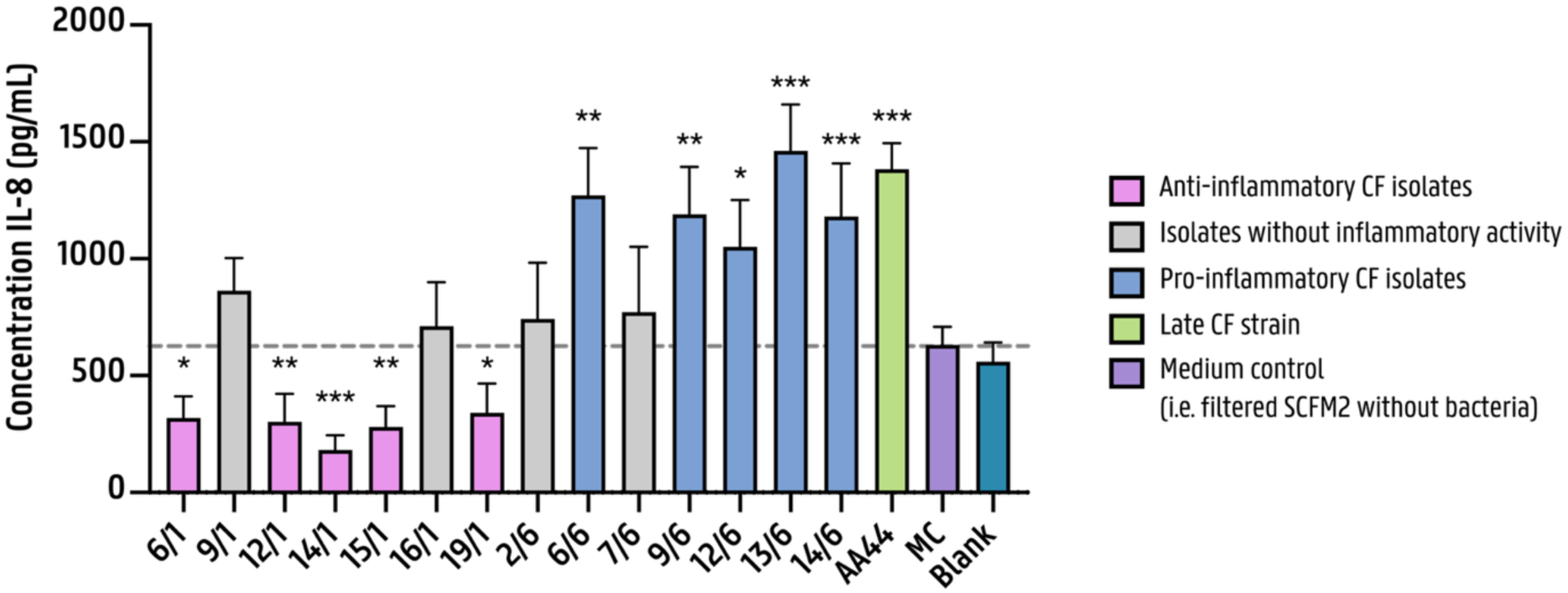
A diverse inflammatory response was observed between isolates, ranging from undetectable to significant levels of IL-8 secretion by 3D lung epithelial cells: Cytokine release was quantified in organotypic 3D lung aggregates following exposure to 40% (v/v) P. aeruginosa cell-free supernatants for 4 h. SCFM2 medium alone served as the medium control (MC), while the blank represents untreated A549 cells cultured in GTSF-2 medium without FBS and antibiotics. Both blank and MC indicate the basal level of IL-8 produced in the absence of a pro-inflammatory stimulus. The positive control was 10% (v/v) supernatant of P. aeruginosa AA44 grown in SCFM2. IL-8 levels were determined using ELISA with cytokine concentration displayed on the y-axis. (n=3–7, statistical analysis using a Tweedie generalized linear mixed model was conducted on the absolute data, which was then compared to the MC. * p < 0.05, ** p < 0.01, *** p < 0.001, error bars represent the standard error). Data previously published in (16).

**Figure S.2:**
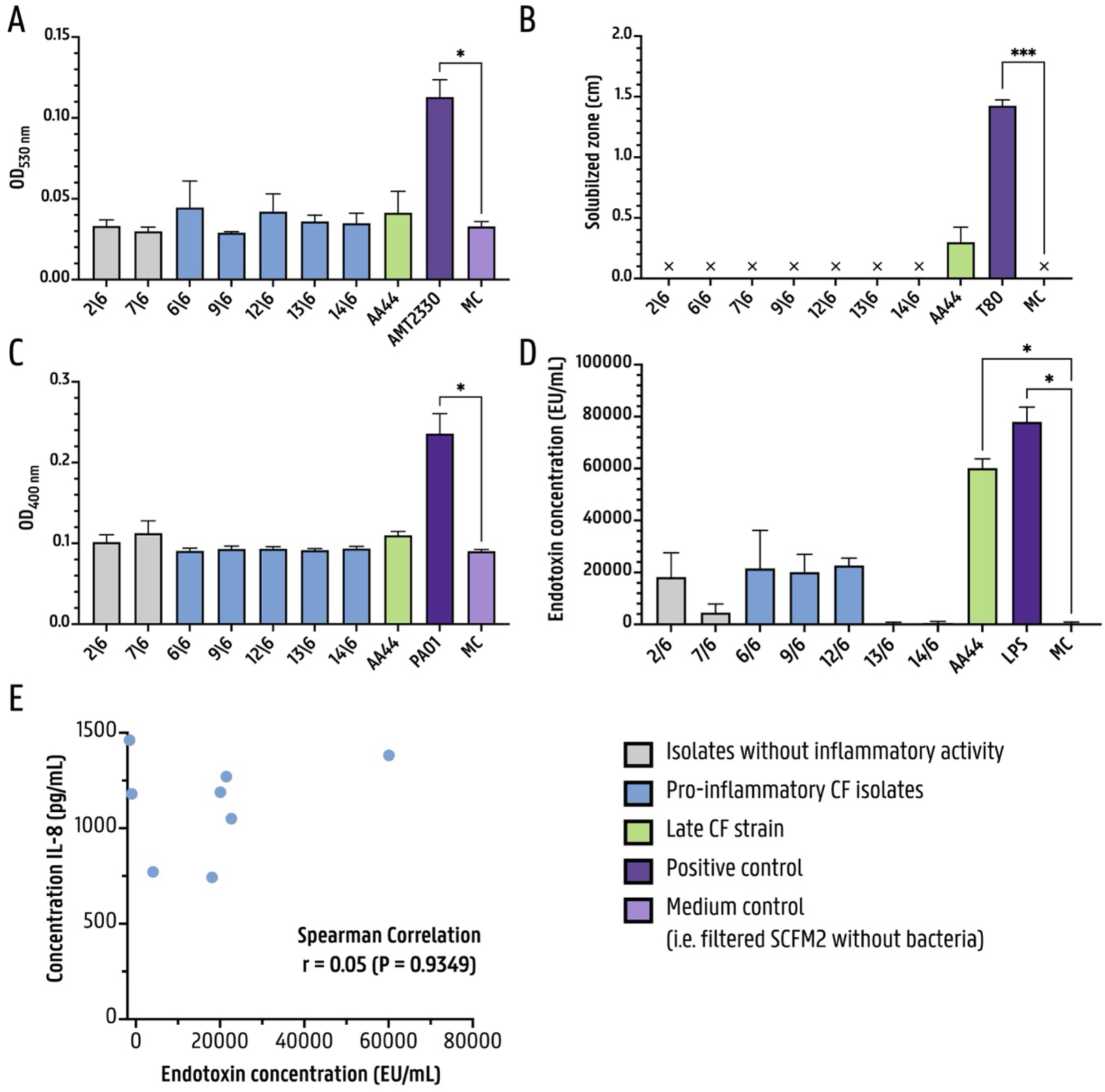
Virulence factors quantified in P. aeruginosa cell-free supernatants or bacterial suspension. (5 x 10^7^ CFU/mL for rhamnolipid assay): (A) Pyocyanin was quantified using chloroform-HCl extraction. Strain AMT 0023-30 served as a positive control. (B) Rhamnolipids were assessed by plating bacterial suspensions on CTAB-MB plates for 48 h under microaerophilic conditions. The size of the halos minus the diameter of the cut-out holes indicated rhamnolipid production. Tween-80 (100 mM) was used as a positive control. An ‘X’ on the x-axis indicates that no measurable halo was present. (C) Pyoverdine levels were measured via absorbance at 400 nm. Strain PAO1 served as a positive control. (D) Endotoxin levels were quantified using the Pierce Chromogenic Endotoxin Quant Kit. LPS from P. aeruginosa serotype 10 (1 mg/mL) was used as a positive control. Although CF isolates 6/6, 9/6, and 12/6 reached endotoxin levels of ∼20,000 EU/mL, these levels were not significantly different from the SCFM2 control (i.e., p = 0.193, 0.176, 0.141, respectively). (E) Scatter plot illustrating the relationship between endotoxin levels in the cell-free supernatants of clinical CF isolates (x-axis), and IL-8 release from 3D lung epithelial aggregates (y-axis) following exposure to 40% (v/v) of the corresponding cell-free supernatants for 4 h. (For assays A-D: n ≥ 3, statistical analysis was performed using the non-parametric Kruskal– Wallis ANOVA comparing to the MC, followed by the Benjamini–Hochberg correction (FDR of 5%). For assay E: n = 8, statistical analysis was performed using a two-tailed Spearman correlation, * p < 0.05, ** p < 0.01, *** p < 0.001, error bars represent standard error)

**Figure S.3:**
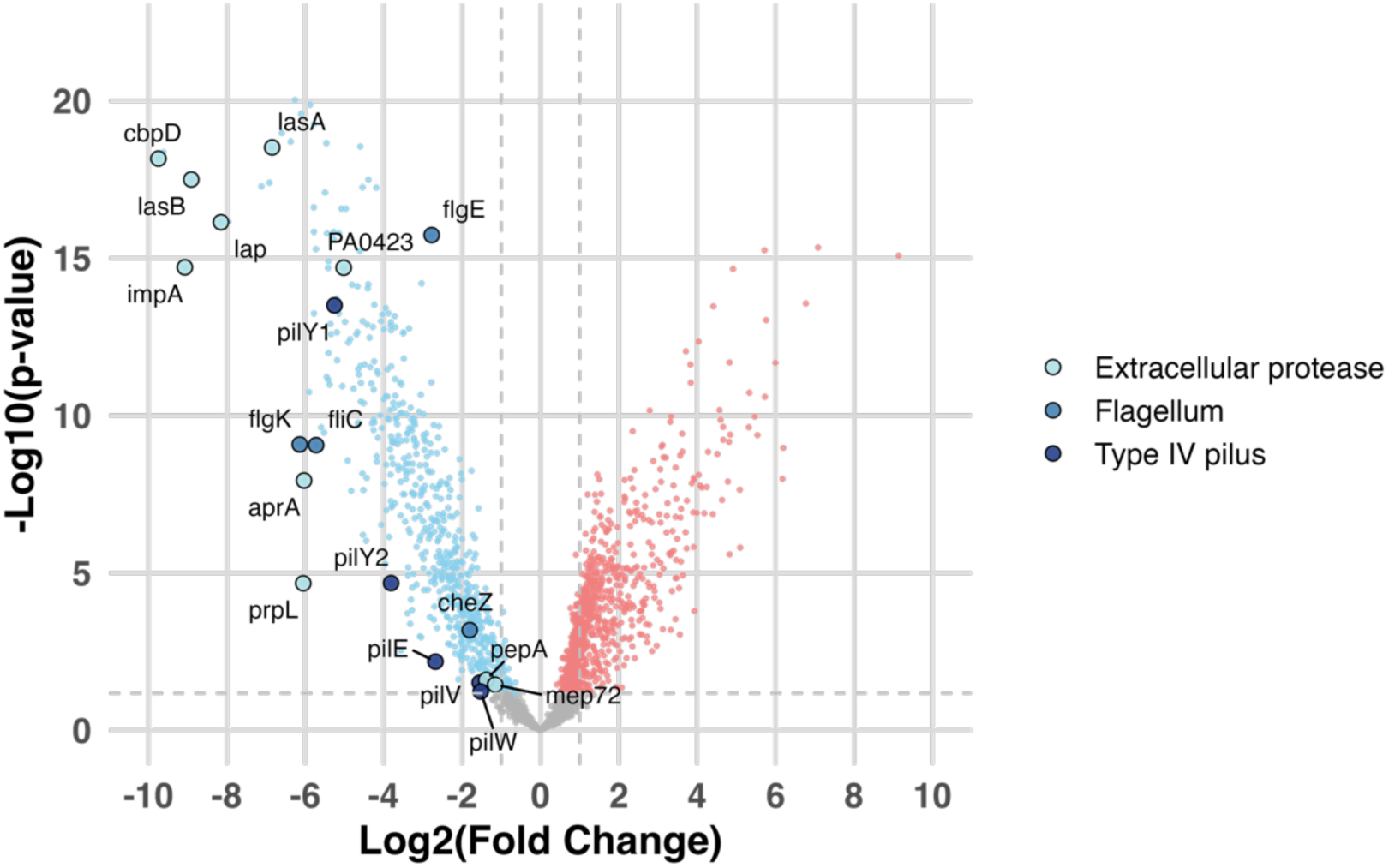
Extracellular proteases and proteins associated with the flagellum or type IV pilus are downregulated in the pro-inflammatory P. aeruginosa isolate group: A volcano plot was generated to display the differences in protein abundance between the pro- and anti-inflammatory isolate group. The difference is represented as log_2_(Fold Change) on the x-axis, plotted against -log_10_(p-values) on the y-axis, derived from a two-sample t-test. Symbols corresponding to extracellular proteases, and proteins associated with the flagellum or type IV pilus are enlarged and labeled. The horizontal dashed line indicates the significance threshold, determined using a permutation-based FDR correction of 5%, and the vertical dashed line marks a log_2_(Fold Change) = |1|.

**Figure S.4:**
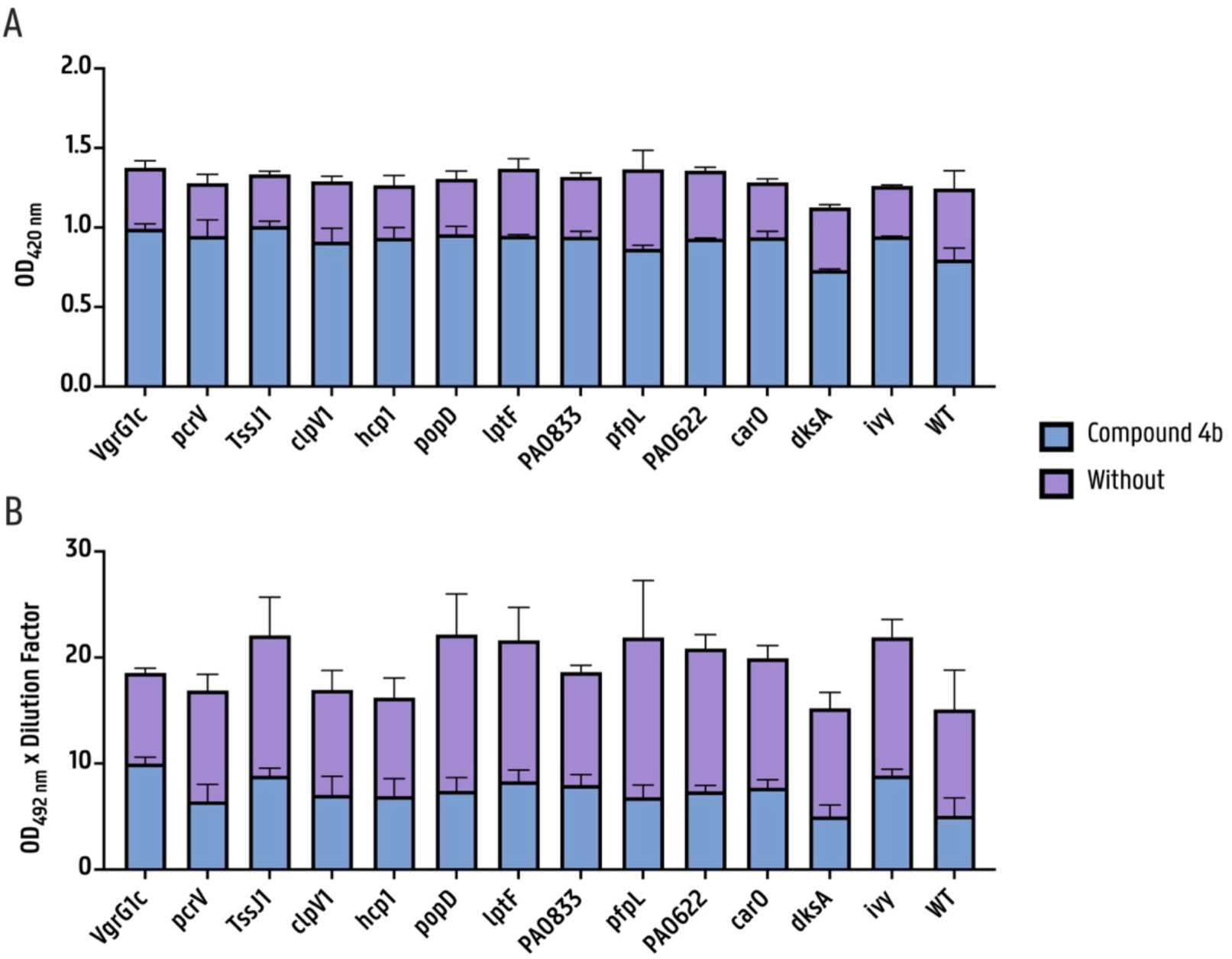
Proteolytic activity in the cell-free culture supernatant of P. aeruginosa PA14 Tn mutant strains: (A) Proteolytic activity of the cell-free supernatants (40% (v/v)) from the PA14 Tn mutant strains was determined by the azocasein colorimetric assay, in the presence or absence of 50 μM LasB inhibitor **4b** (0.5% (v/v)). (B) Elastolytic activity of the cell-free supernatants (40% (v/v)) from the PA14 Tn mutant strains was determined by the Elastin-Congo red assay, in the presence or absence of 50 μM compound **4b** (0.5% (v/v)). High absorbance values correspond with high proteolytic/elastolytic activity. (For all assays: n ≥ 3, statistical analysis was performed using a mixed-effects model with REML estimation, error bars represent the standard error).

**Figure S.5:**
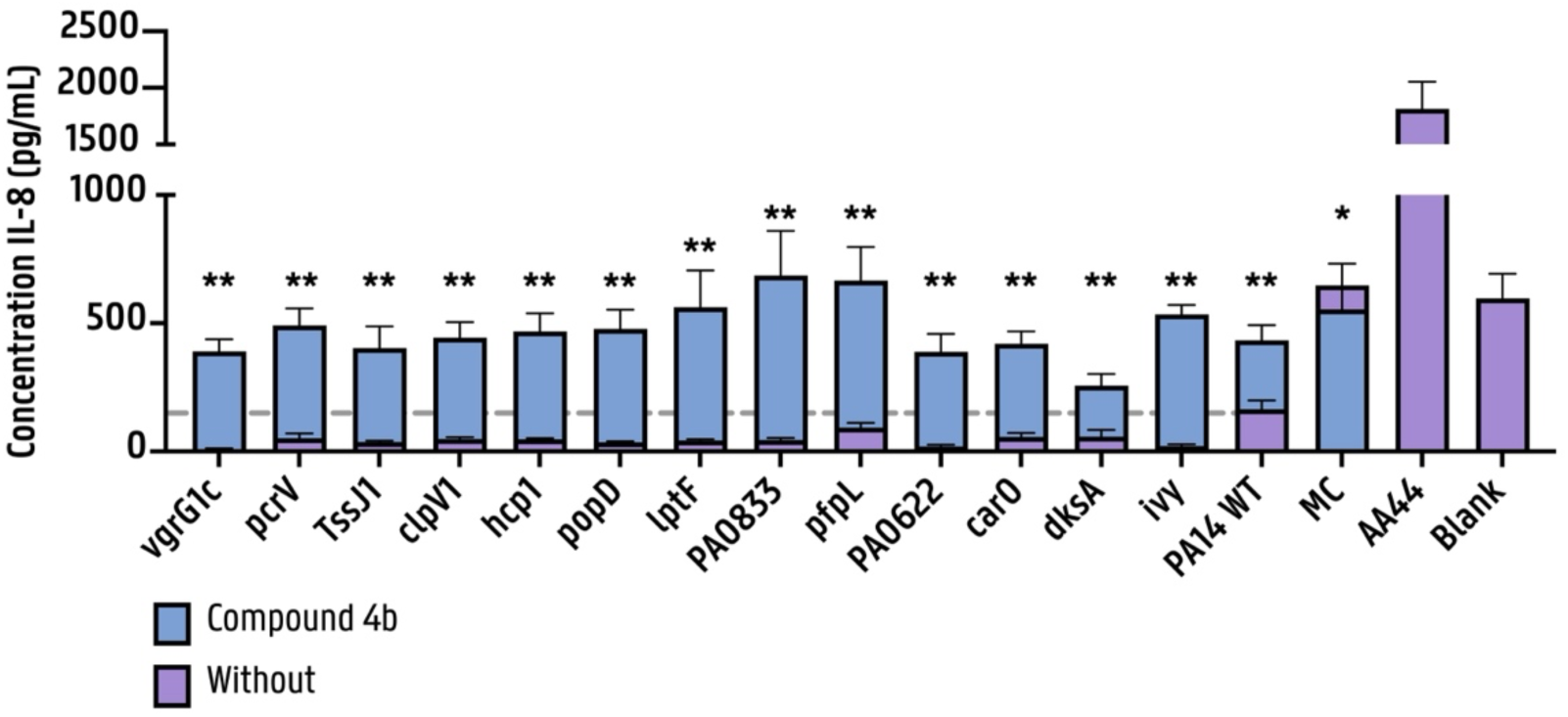
IL-8 produced by 3D lung aggregates following exposure to cell-free supernatants from P. aeruginosa PA14 Tn mutant strains: IL-8 release was measured in an organotypic 3D lung cell culture model after 4 h exposure to 40% (v/v) P. aeruginosa cell-free supernatants, in the presence or absence of 50 μM LasB inhibitor **4b** (0.5% (v/v)). Differences in IL-8 release in the presence or absence of compound **4b** were assessed for each Tn mutant. SCFM2 alone served as the medium control (MC), while the blank represents untreated A549 cells cultured in GTSF-2 medium without FBS. Both controls reflect basal cytokine levels in the absence of a pro-inflammatory stimulus. (n ≥ 3, statistical analysis was performed using multiple Mann-Whitney U tests followed by the Benjamini-Hochberg correction (FDR = 5%) for multiple testing to assess the effect of the inhibitor for each pair (asterisk above bars indicate significant differences). Additionally, the non-parametric Kruskal–Wallis ANOVA followed by Benjamini–Hochberg correction (5% FDR) was used to compare IL-8 release (in the absence of inhibitor **4b**) induced by each Tn mutant with that of the PA14 WT supernatant, no significant differences were observed (not shown), * p < 0.05, ** p < 0.01, *** p < 0.001, error bars represent the standard error).

**Figure S.6:**
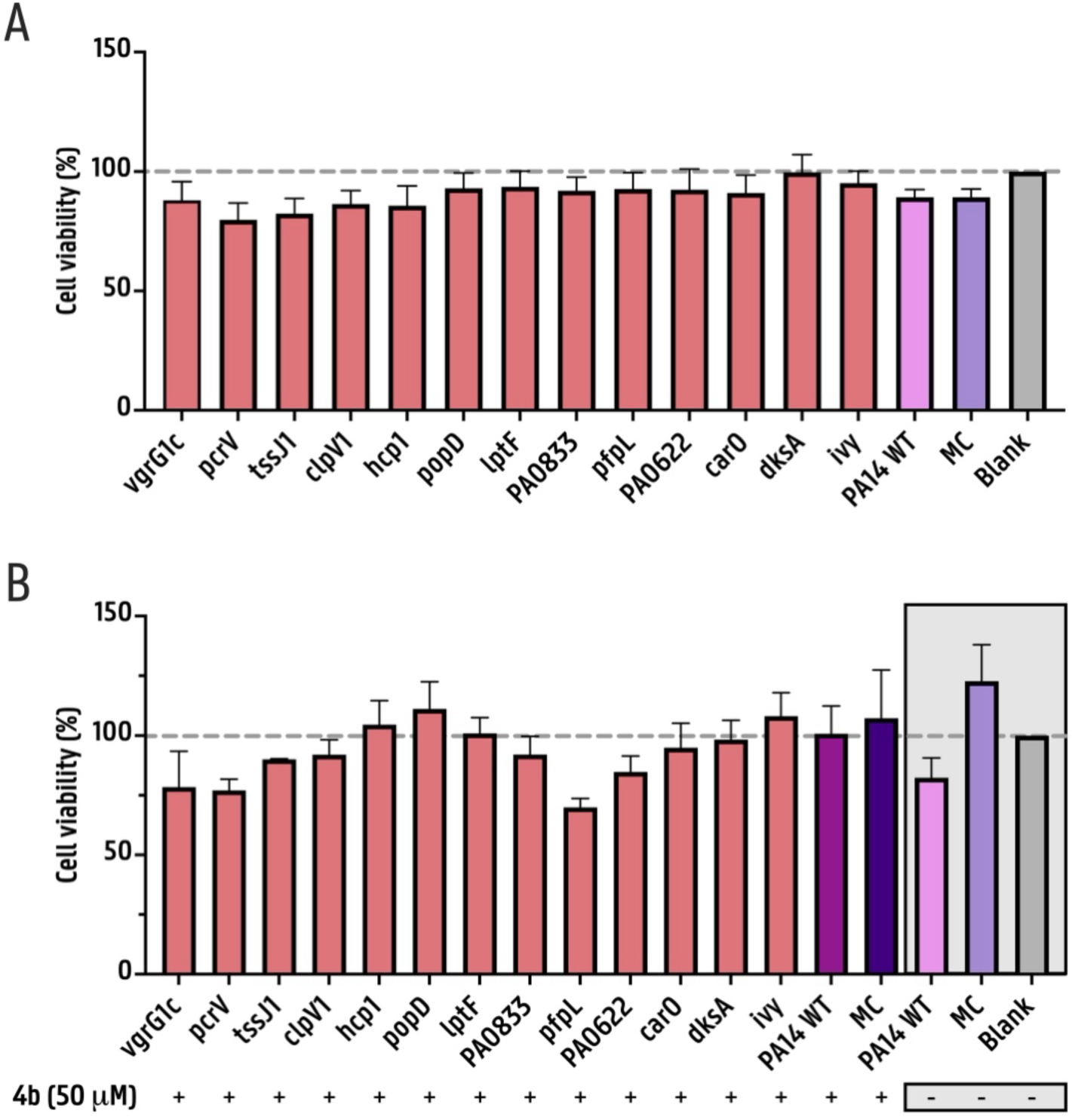
Cytotoxic response of 3D lung aggregates to cell-free supernatants from PA14 Tn mutant strains, with or without LasB inhibitor 4b: (A) Cell-viability of the organotypic 3D lung cell culture model following 4h exposure to 40% (v/v) P. aeruginosa cell-free supernatants. (B) Cell-viability of the organotypic 3D lung cell culture model after 4h exposure to 40% (v/v) P. aeruginosa cell-free supernatants, in the presence or absence of 50 μM LasB inhibitor 4b (0.5% (v/v)). Viability was determined by LDH release using an intracellular LDH assay. The blank control represents untreated A549 cells in GTSF-2 medium without FBS, while SCFM2 alone was used as a medium control (MC). The y-axis shows the cell-viability relative to the blank. (For all assays: n ≥ 3, statistical analysis of the relative data was conducted using the non-parametric Kruskal– Wallis ANOVA as described in (37), multiple testing correction was performed using the Benjamini–Hochberg procedure with a 5% FDR, no significant differences were observed (not shown), * p < 0.05, error bars represent the standard error).

**Figure S.7:**
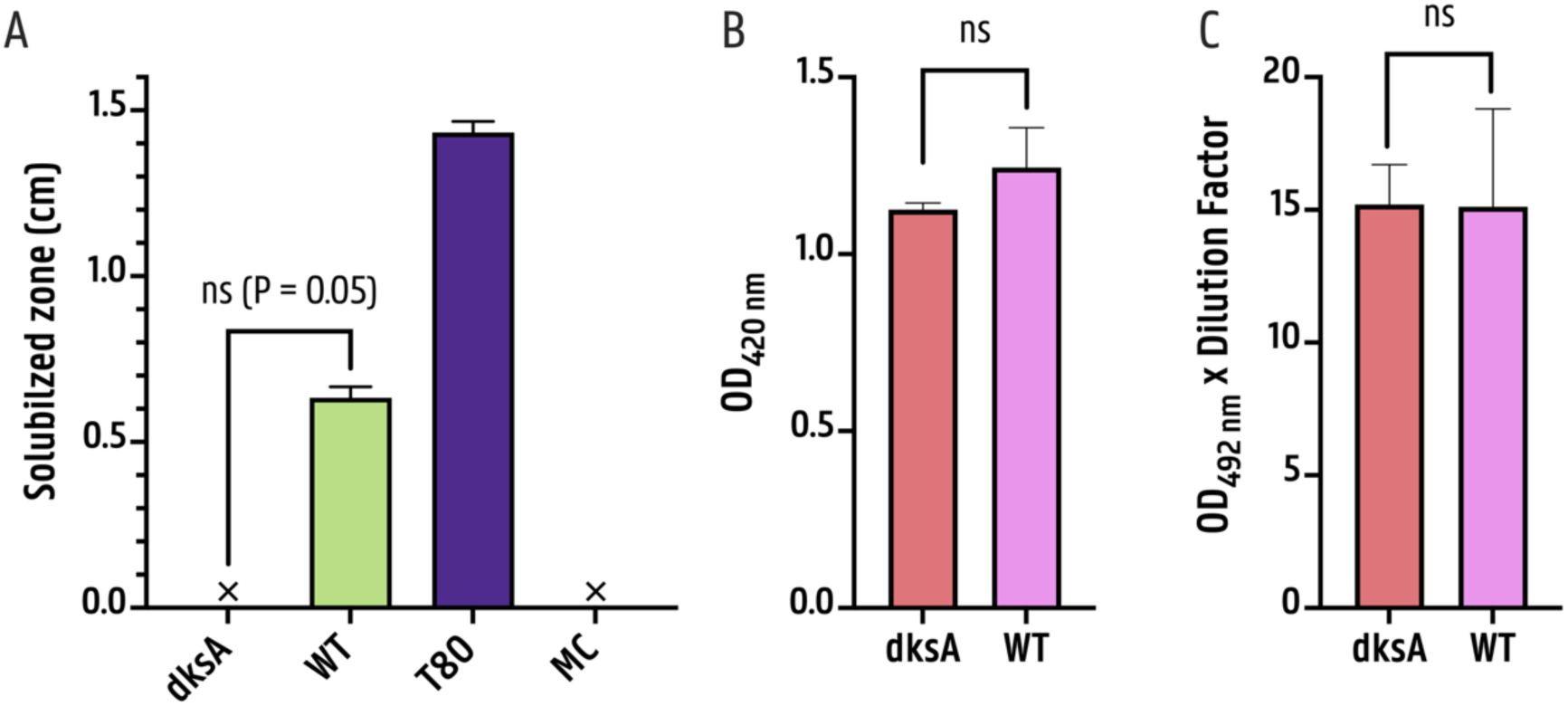
Virulence factors were quantified in PA14 WT and PA14 dksA::Tn cell-free supernatant or bacterial suspension of PA14 WT and PA14 dksA::Tn. ( x 10^7^ CFU/mL for rhamnolipid assay): (A) Rhamnolipids were assessed by plating bacterial suspensions on CTAB-MB plates for 48 h under microaerophilic conditions. The size of the halos minus the diameter of the cut-out wells indicated rhamnolipid production. Tween-80 was used as a positive control. An ‘X’ on the x-axis indicates that no measurable halo was present. (B) Proteolytic activity of PA14 WT and PA14 dksA::Tn cell-free supernatant was determined by the azocasein colorimetric assay. (C) Elastolytic activity of PA14 WT and PA14 dksA::Tn cell-free supernatant was determined by the Elastin-Congo red assay. (For all assays: n = 3, statistical analysis was performed using a one-tailed non-parametric Mann–Whitney U test comparing PA14 dksA::Tn to PA14 WT, * p < 0.05, error bars represent standard error).

**Figure S.8:**
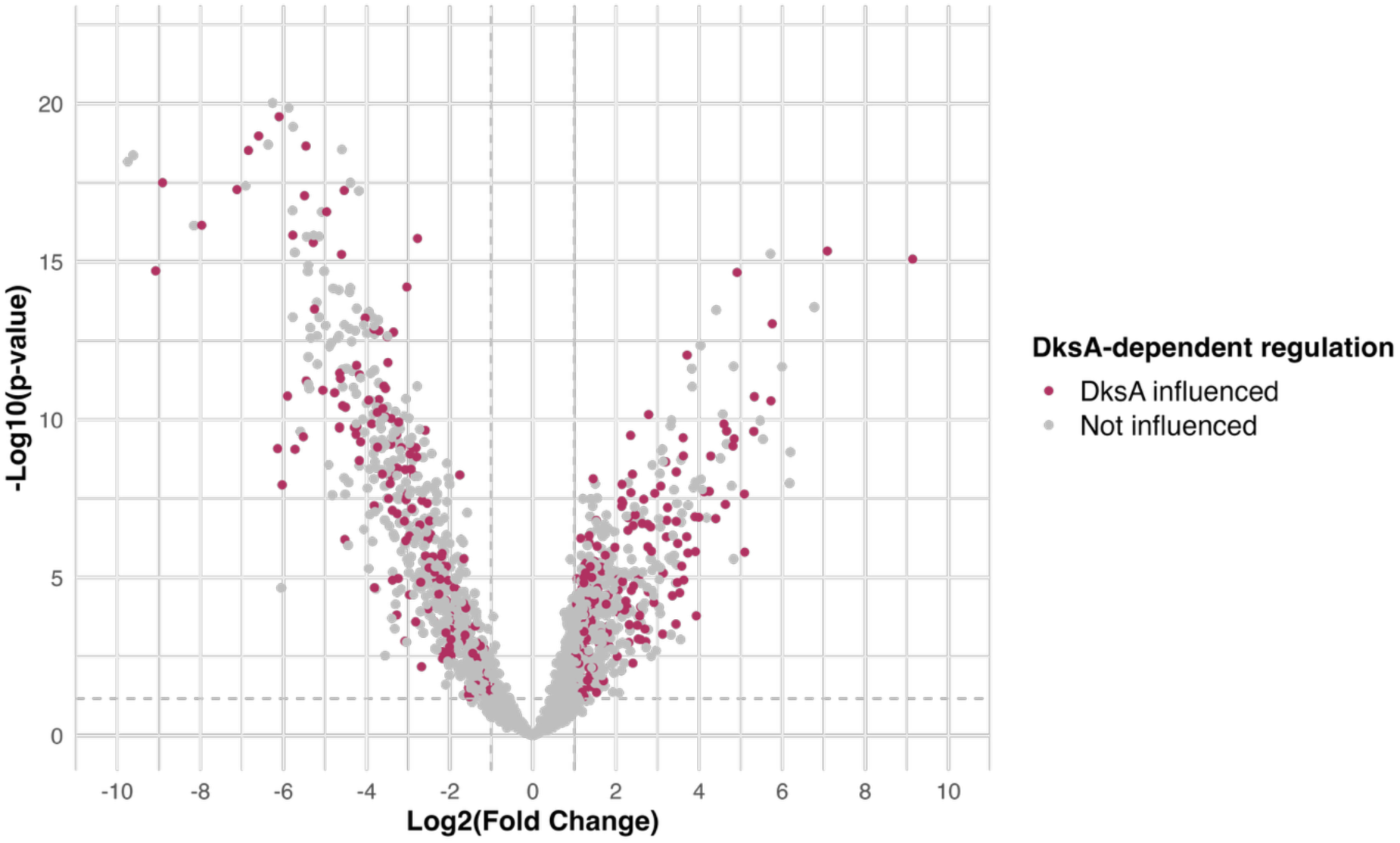
dksA-dependent regulation of differentially expressed proteins in pro-versus anti-inflammatory isolates: Volcano plot showing protein abundance differences between pro-inflammatory and anti-inflammatory isolates. Differences are represented as log_2_(Fold Change) on the x-axis, against -log_10_(p-values) on the y-axis, calculated using a two-sample t-test. Proteins significantly differentially expressed with a fold change > |2| under dksA regulation, based on data from Fortuna et al. (52), are highlighted. The horizontal dashed line indicates the significance threshold (5% permutation-based FDR), and the vertical dashed line marks a log_2_(Fold Change) = |1|.

**Figure S.3:**
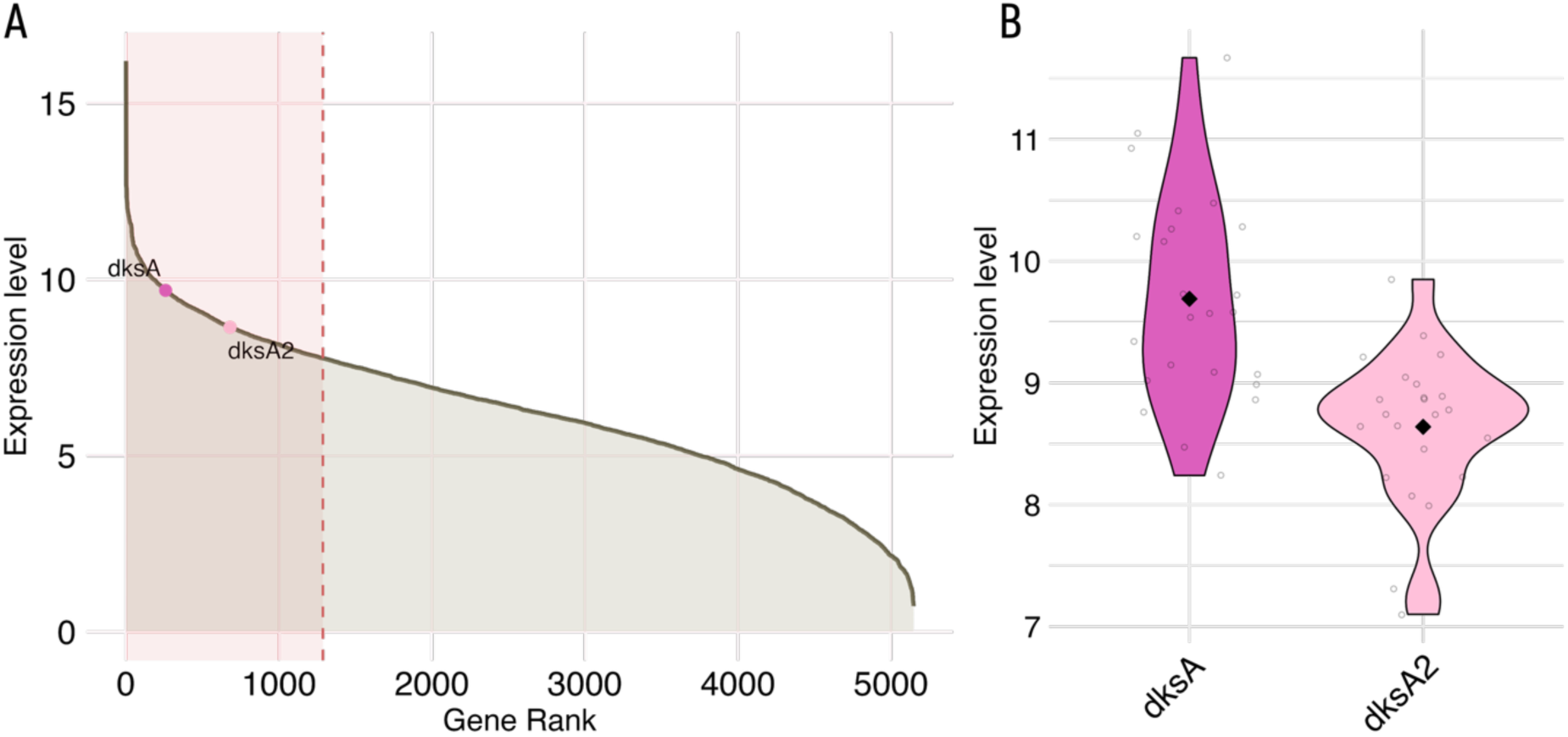
Expression profile of dksA and dksA2 across 24 P. aeruginosa transcriptomes derived from CF sputum samples (data from Lewin et al. (43)): (A) Cumulative gene expression plot displaying genes ranked by their average expression (VST-normalized count data) across the 24 CF sputum-derived P. aeruginosa transcriptomes. dksA and dksA2 are labeled and the top quartile (25%) of most highly expressed genes is highlighted with a transparent red box; (B) Violin plots depicting the distribution and variability of expression levels for dksA and dksA2 across all 24 samples with each violin representing one gene.

**Figure S.10:**
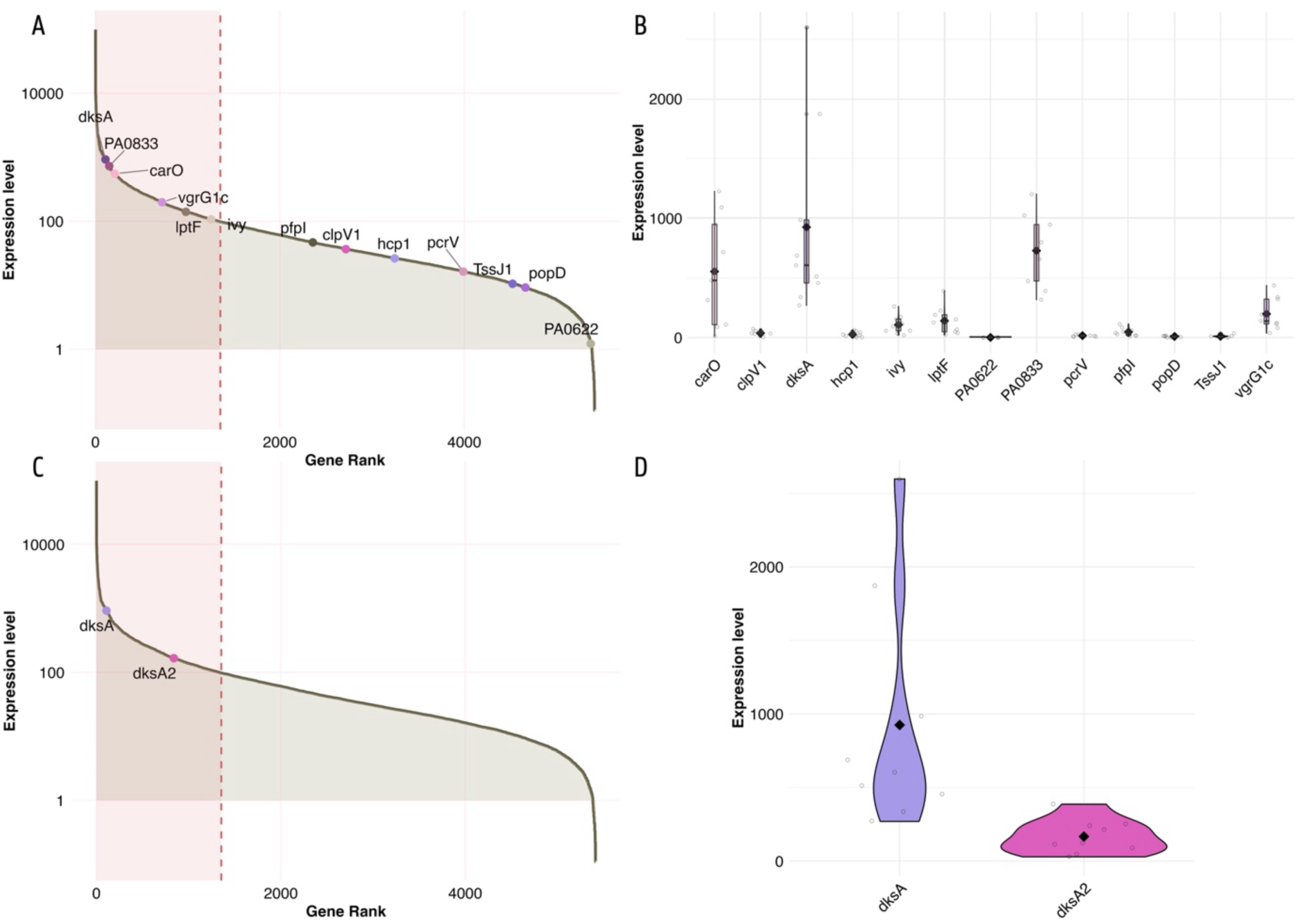
Expression profile of mediators of interest across nine P. aeruginosa transcriptomes derived from CF sputum samples belonging to four different pwCF chronically infected with P. aeruginosa: data were obtained from the P. aeruginosa database (originally derived from Rossi et al. (44)) (A) Cumulative gene expression plot displaying genes ranked by their average expression (normalized count data) across the nine CF sputum-derived P. aeruginosa transcriptomes. Proteins of interest are labeled, and the top quartile (25%) of most highly expressed genes is highlighted with a transparent red box; (B) Violin plots depicting the distribution and variability of expression levels for selected mediators across all nine samples with each violin representing one gene. (C) Cumulative gene expression plot with dksA and dksA2 labeled. (D) Violin plots depicting the distribution and variability of expression levels for dksA and dksA2 across all nine samples.

**Table S.1:**
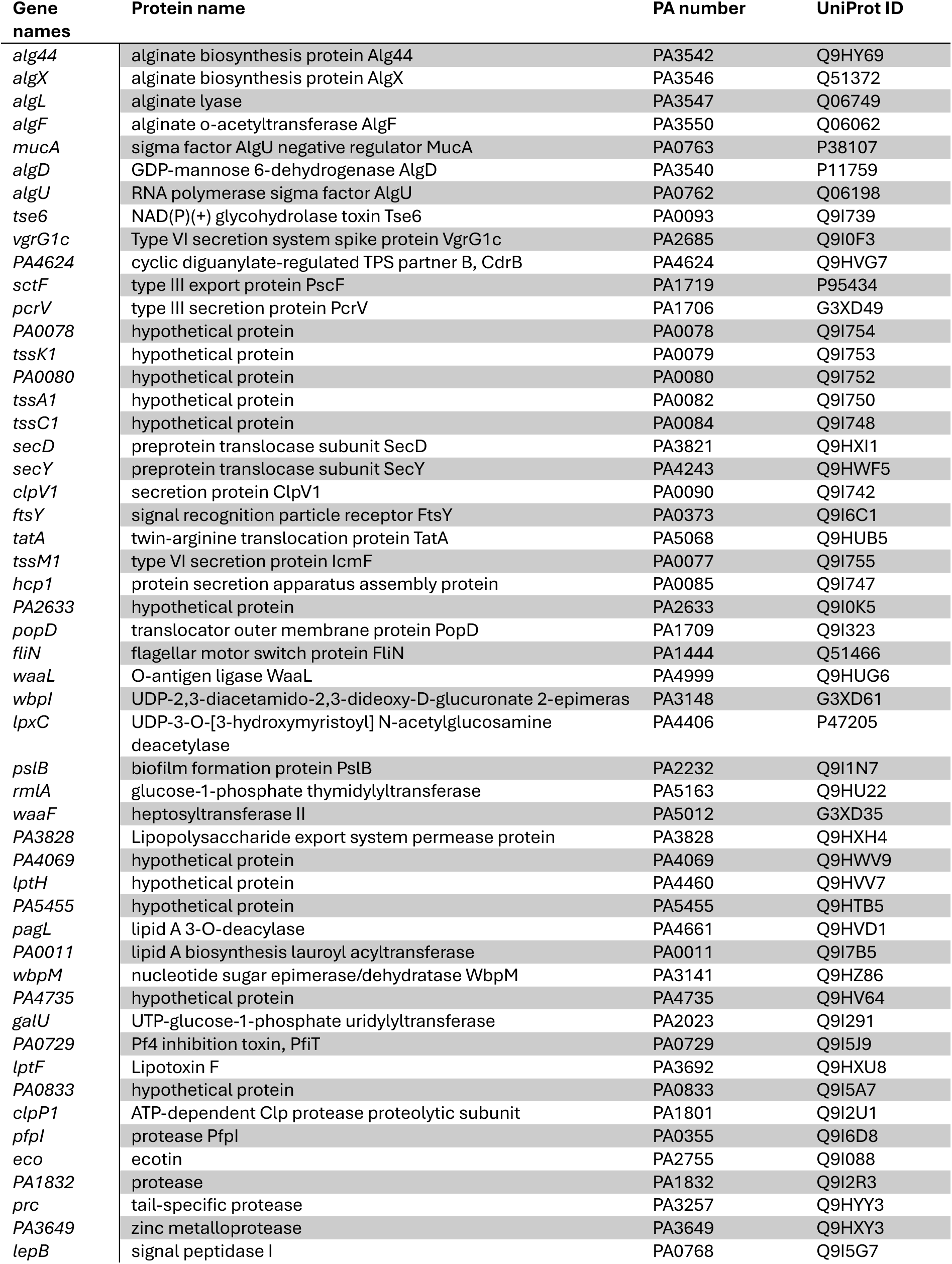
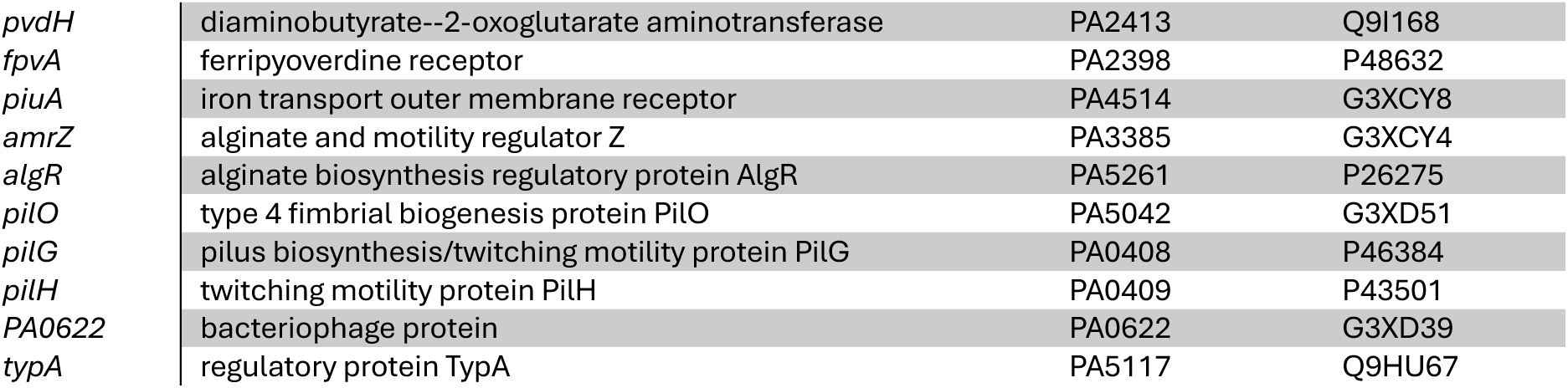
Sixty-two virulence-associated proteins were upregulated in isolates that triggered pro-inflammatory activity in 3D lung aggregates (p < 0.05, log_2_(Fold Change) > 1). The complete dataset of the 62 proteins with DAVID functional annotation is provided in the accompanying Excel file.

**Table S.2:**
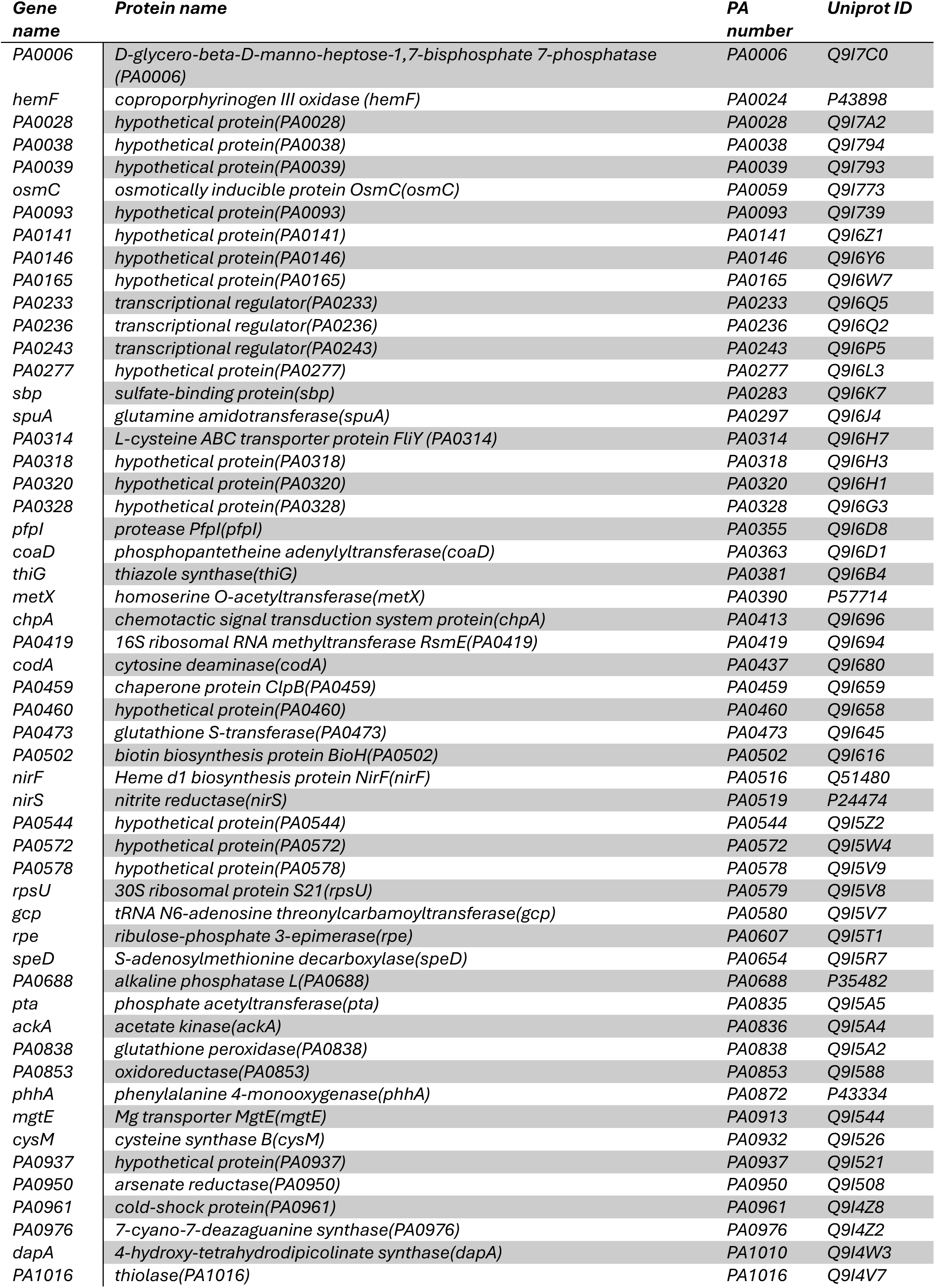
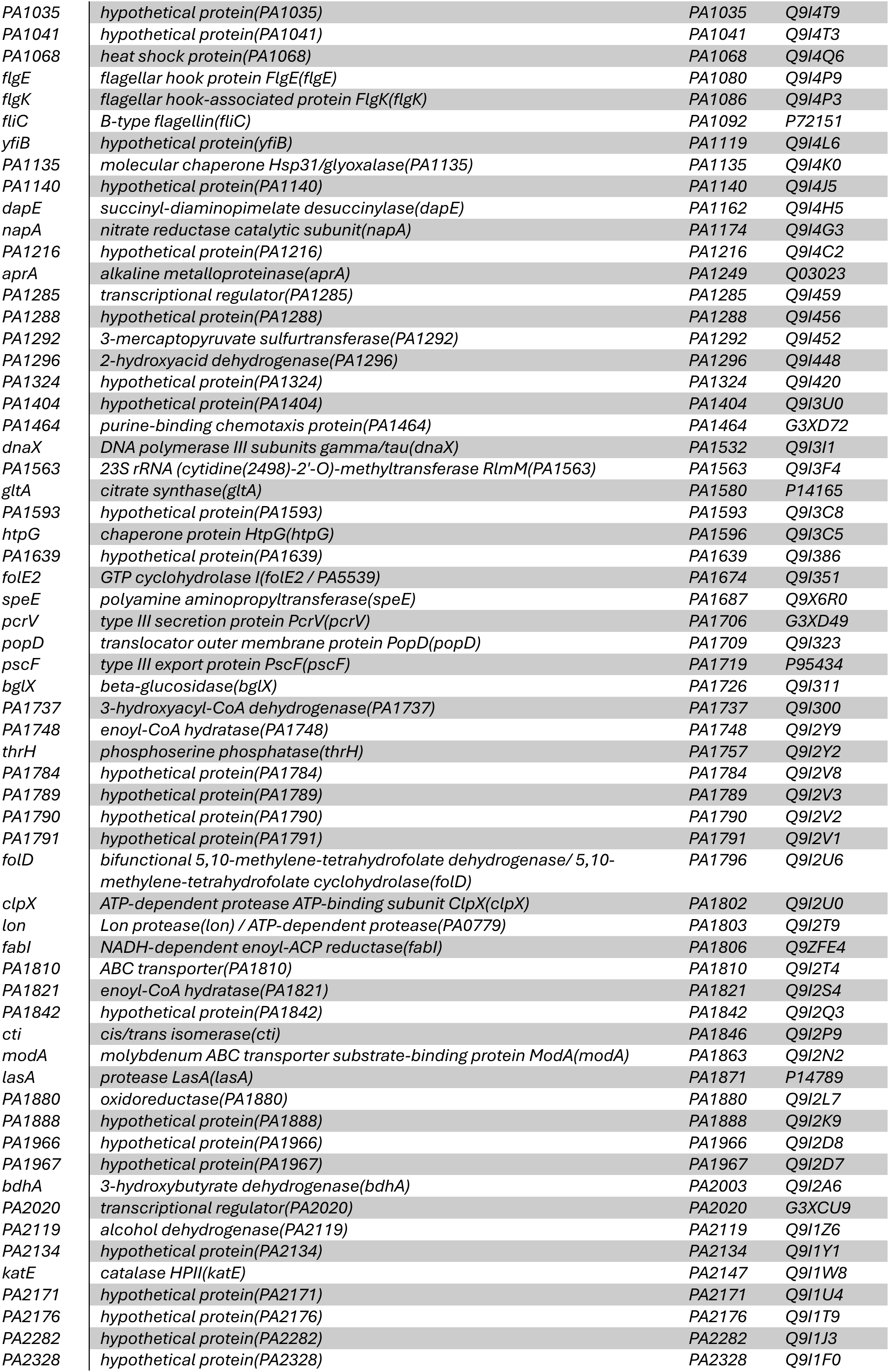
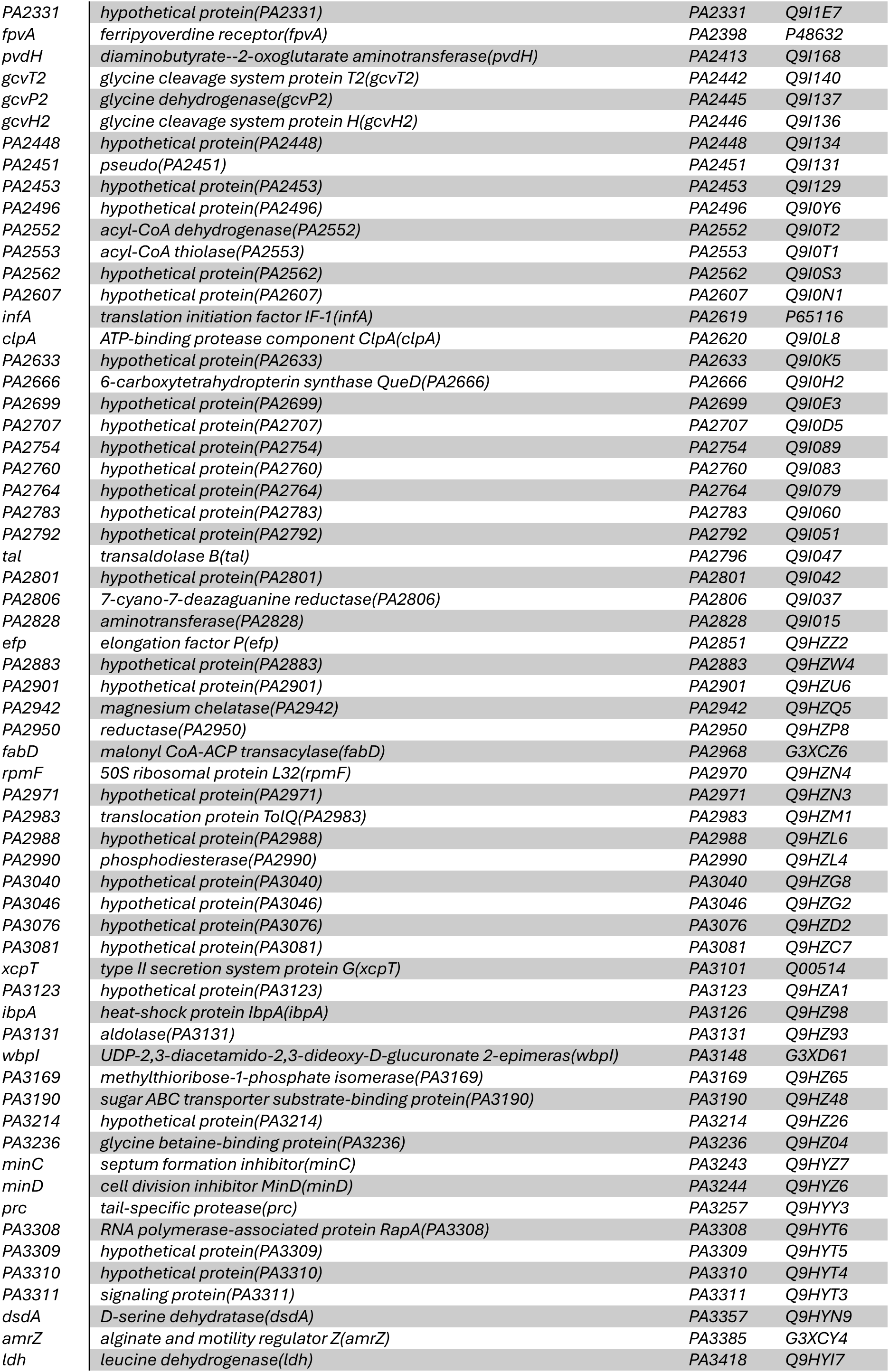
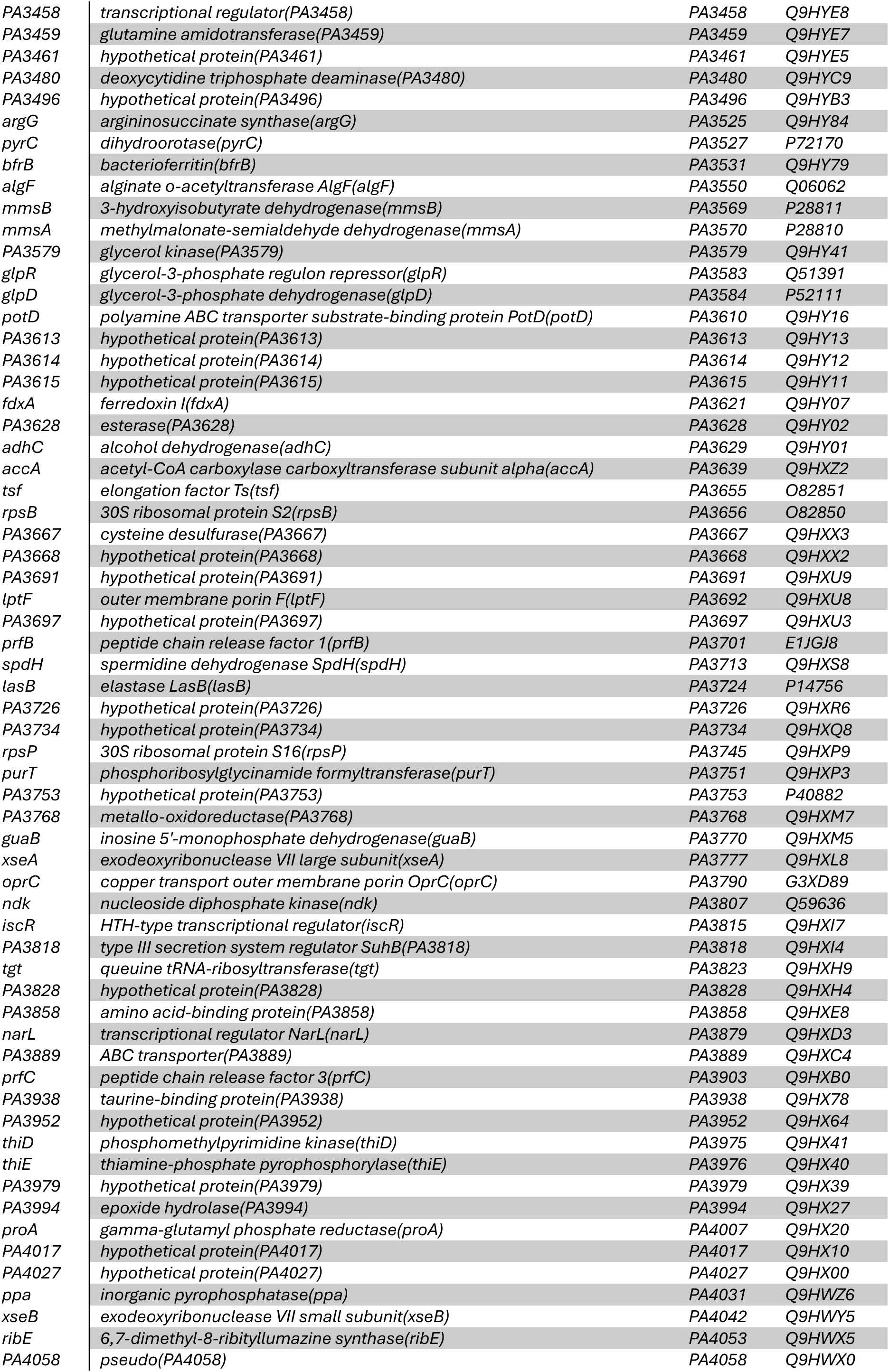
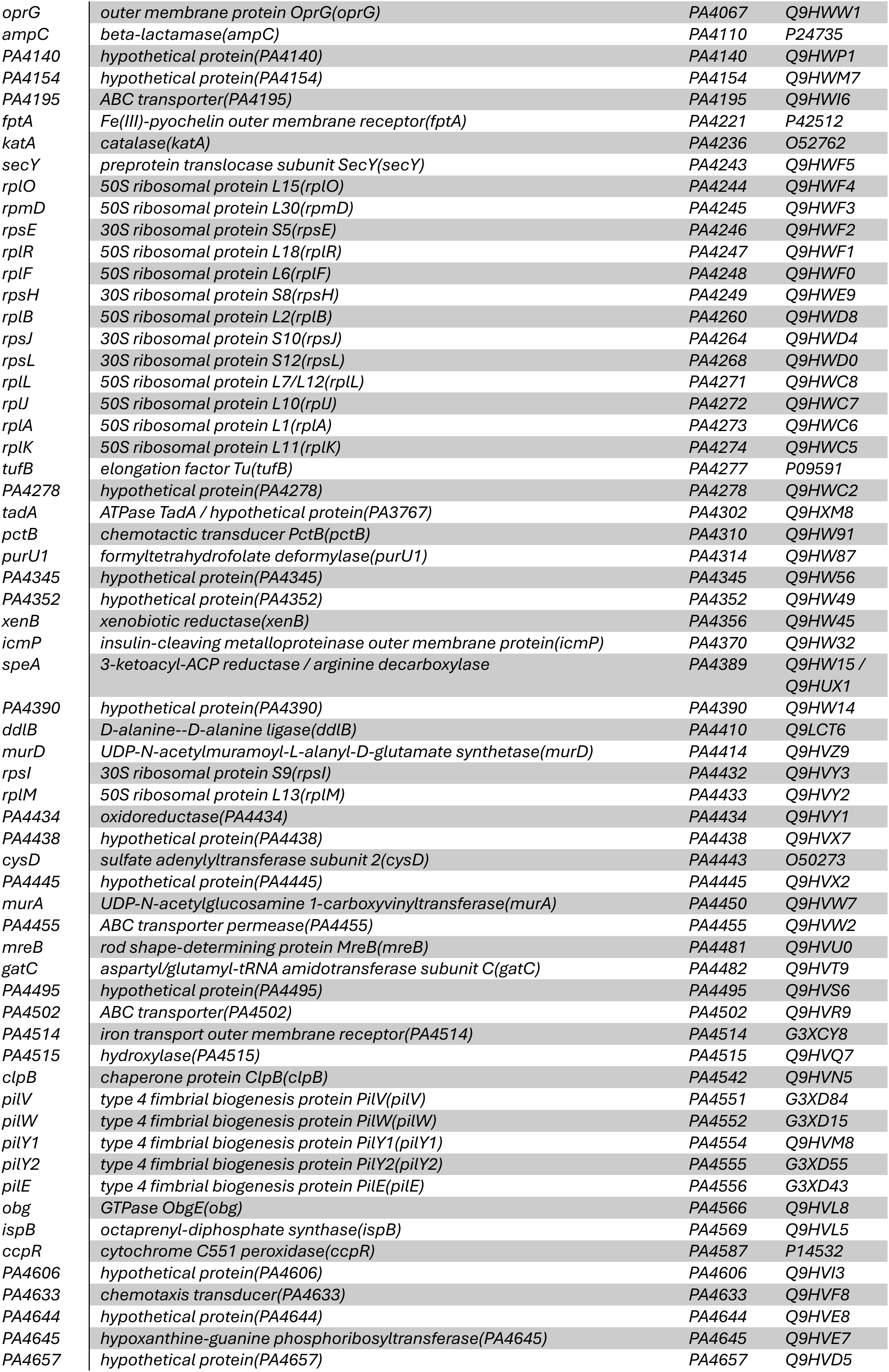
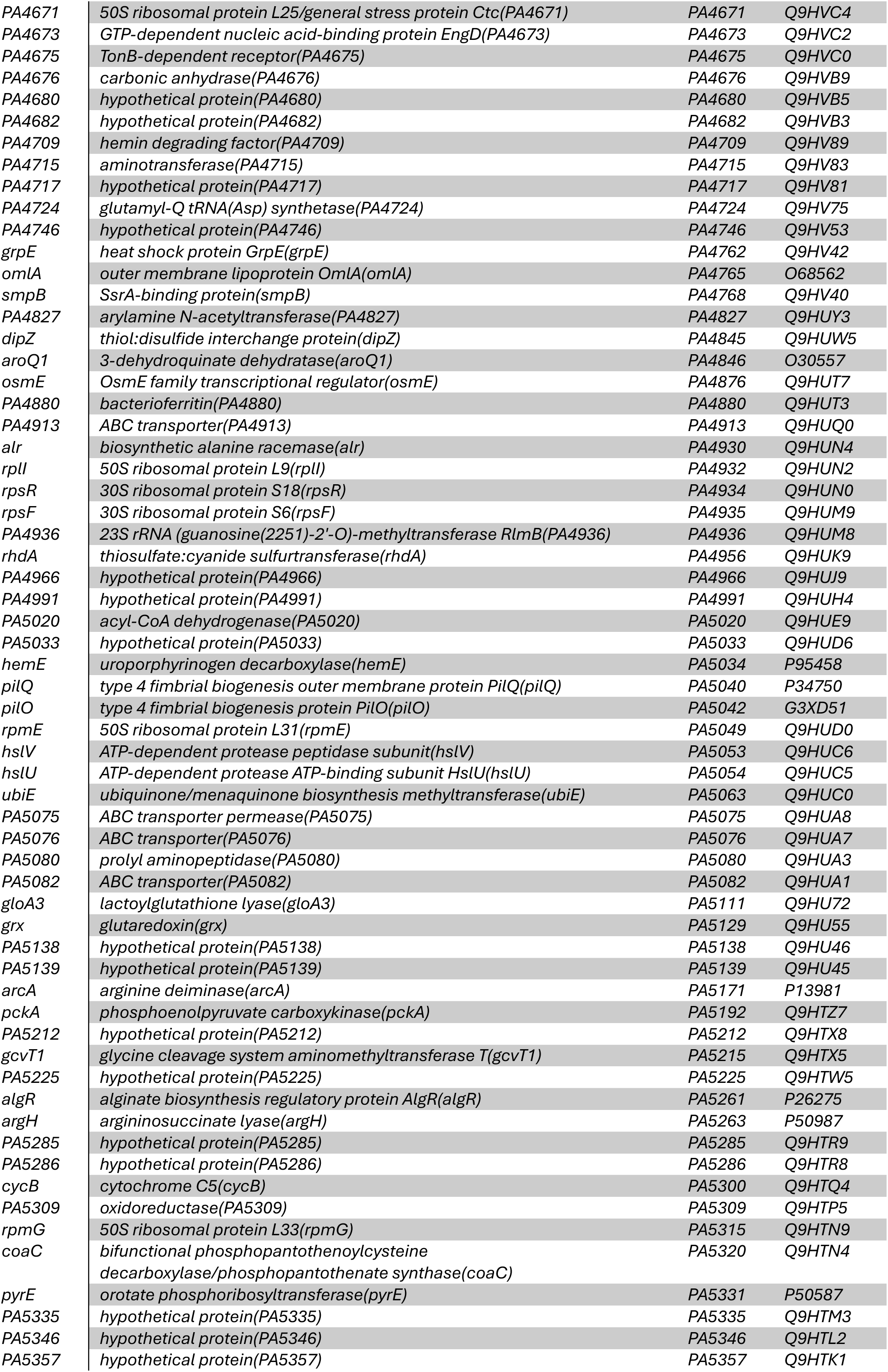
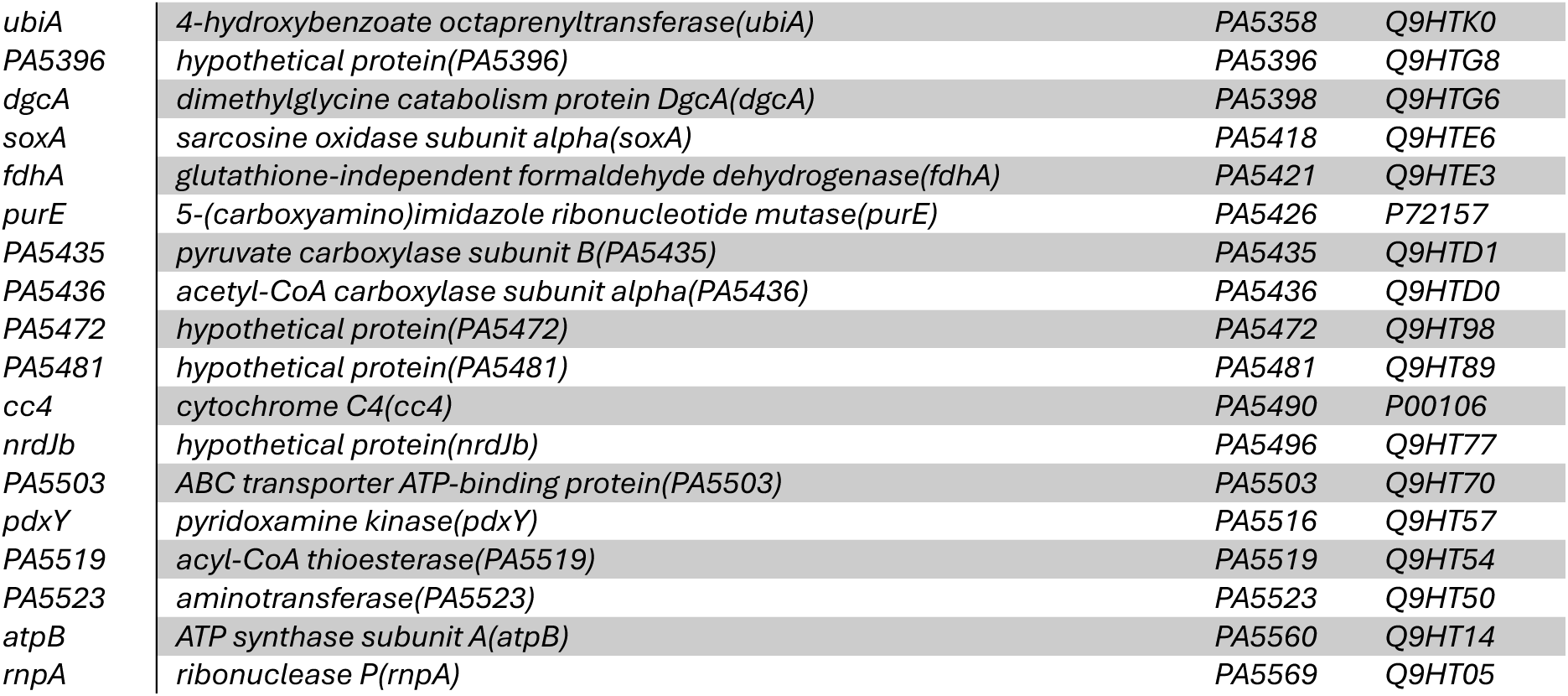
dksA-regulated proteins differentially expressed (p < 0.05, log_2_(Fold Change) > |1|) between pro- and anti-inflammatory isolates: dksA regulon based on data from Fortuna et al. (52).

## References

1. Kerem E, Viviani L, Zolin A, et al. Factors associated with FEV1 decline in cystic fibrosis: analysis of the ECFS Patient Registry. European Respiratory Journal. 2013;43(1):125–33.

2. Robertson JK, Goldberg JB, Robertson JK, et al. The impact of cystic fibrosis transmembrane conductance regulator (CFTR) modulators on the pulmonary microbiota. Microbiology. 2025;171(4):001553.

3. Valladares KN, Jones LI, Barnes JW, et al. Highly Effective Modulator Therapy: Implications for the Microbial Landscape in Cystic Fibrosis. International Journal of Molecular Sciences. 2024;25(22):11865.

4. Nickerson R, Thornton CS, Johnston B, et al. *Pseudomonas aeruginosa* in chronic lung disease: untangling the dysregulated host immune response. Frontiers in Immunology. 2024;15:1405376.

5. Cohen TS, Prince A. Cystic fibrosis: a mucosal immunodeficiency syndrome. Nature Medicine. 2012;18(4):509–19.

6. Letizia M, Diggle SP, Whiteley M. *Pseudomonas aeruginosa*: ecology, evolution, pathogenesis and antimicrobial susceptibility. Nature Reviews Microbiology. 2025;23:701–17.

7. Cystic Fibrosis Foundation. 2024 Patient Registry Highlights. 2024.

8. 8. Zolin A, Adamoli A, Bakkeheim E, et al. European Cystic Fibrosis Society Patient Registry Annual Report 2023. European Cystic Fibrosis Society Patient; 2023.

9. Faure E, Kwong K, Nguyen D. *Pseudomonas aeruginosa* in Chronic Lung Infections: How to Adapt Within the Host? Front Immunol. 2018;9:2416.

10. Rossi E, La Rosa R, Bartell JA, et al. *Pseudomonas aeruginosa* adaptation and evolution in patients with cystic fibrosis. Nature Reviews Microbiology. 2020;19(5):331–42.

11. LaFayette SL, Houle D, Beaudoin T, et al. Cystic fibrosis-adapted *Pseudomonas aeruginosa* quorum sensing *lasR* mutants cause hyperinflammatory responses. Sci Adv. 2015;1(6):e1500199.

12. Bragonzi A, Paroni M, Nonis A, et al. *Pseudomonas aeruginosa* Microevolution during Cystic Fibrosis Lung Infection Establishes Clones with Adapted Virulence. American Journal of Respiratory and Critical Care Medicine. 2009;180(2):138–45.

13. Phuong MS, Hernandez RE, Wolter DJ, et al. Impairment in inflammasome signaling by the chronic *Pseudomonas aeruginosa* isolates from cystic fibrosis patients results in an increase in inflammatory response. Cell Death & Disease. 2021;12(241).

14. Hennemann LC, LaFayette SL, Malet JK, et al. LasR-deficient *Pseudomonas aeruginosa* variants increase airway epithelial mICAM-1 expression and enhance neutrophilic lung inflammation. PLoS Pathog. 2021;17(3):e1009375.

15. Rossi E, Lausen M, Øbro NF, et al. Widespread alterations in systemic immune profile are linked to lung function heterogeneity and airway microbes in cystic fibrosis. Journal of Cystic Fibrosis. 2024;23(5):885–95.

16. Wauters M, Van Den Bossche S, Grassi L, et al. Unraveling the immunosuppressive role of Elastase B produced by cystic fibrosis isolates of *Pseudomonas aeruginosa* in an organotypic 3D lung epithelial cell model. Virulence. 2025;16(1).

17. Turner KH, Wessel AK, Palmer GC, et al. Essential genome of *Pseudomonas aeruginosa* in cystic fibrosis sputum. Proc Natl Acad Sci U S A. 2015;112(13):4110–5.

18. Cornforth DM, Diggle FL, Melvin JA, et al. Quantitative Framework for Model Evaluation in Microbiology Research Using *Pseudomonas aeruginosa* and Cystic Fibrosis Infection as a Test Case. Mbio. 2020;11(1):e03042–19.

19. Barrila J, Radtke AL, Crabbe A, et al. Organotypic 3D cell culture models: using the rotating wall vessel to study host-pathogen interactions. Nat Rev Microbiol. 2010;8(11):791–801.

20. Crabbe A, Liu Y, Matthijs N, et al. Antimicrobial efficacy against *Pseudomonas aeruginosa* biofilm formation in a three-dimensional lung epithelial model and the influence of fetal bovine serum. Sci Rep. 2017;7:43321.

21. Carterson AJ, Honer zu Bentrup K, Ott CM, et al. A549 lung epithelial cells grown as three-dimensional aggregates: alternative tissue culture model for *Pseudomonas aeruginosa* pathogenesis. Infect Immun. 2005;73(2):1129-40.

22. Van Den Bossche S, Abatih E, Grassi L, et al. Pooling isolates to address the diversity in antimicrobial susceptibility of *Pseudomonas aeruginosa* in cystic fibrosis. Microbiology Spectrum. 2023;11(6):e0044923.

23. Cullen L, Weiser R, Olszak T, et al. Phenotypic characterization of an international *Pseudomonas aeruginosa* reference panel: strains of cystic fibrosis (CF) origin show less *in vivo* virulence than non-CF strains. Microbiology. 2015;161(10):1961–77.

24. Jacobs MA, Alwood A, Thaipisuttikul I, et al. Comprehensive transposon mutant library of *Pseudomonas aeruginosa*. Proceedings of the National Academy of Sciences. 2003;100(24):14339–44.

25. Chandler CE, Horspool AM, Hill PJ, et al. Genomic and Phenotypic Diversity among Ten Laboratory Isolates of *Pseudomonas aeruginosa* PAO1. Journal of Bacteriology. 2018;201(5):10.1128/jb.00595-18.

26. Liberati NT, Urbach JM, Miyata S, et al. An ordered, nonredundant library of *Pseudomonas aeruginosa* strain PA14 transposon insertion mutants. Proceedings of the National Academy of Sciences of the United States of America. 2006;103(8):2833–8.

27. Liberati NT, Urbach JM, Miyata S, et al. PA14 Non-Redundant Transposon Insertion Mutant Set (PA14NR Set) 2006 [Available from: https://pa14.mgh.harvard.edu/cgi-bin/pa14/home.cgi.

28. Worlitzsch D, Tarran R, Ulrich M, et al. Effects of reduced mucus oxygen concentration in airway Pseudomonas infections of cystic fibrosis patients. The Journal of Clinical Investigation. 2002;109(3):317–25.

29. Konstantinović J, Kany AM, Alhayek A, et al. Inhibitors of the Elastase LasB for the Treatment of *Pseudomonas aeruginosa* Lung Infections. ACS Central Science. 2023;9(12):2205– 15.

30. Zhang Y, Sass A, Van Acker H, et al. Coumarin Reduces Virulence and Biofilm Formation in *Pseudomonas aeruginosa* by Affecting Quorum Sensing, Type III Secretion and C-di-GMP Levels. Front Microbiol. 2018;9:1952.

31. Jiang J, Jin M, Li X, et al. Recent progress and trends in the analysis and identification of rhamnolipids. Appl Microbiol Biotechnol. 2020;104(19):8171–86.

32. Pinzon NM, Ju LK. Improved detection of rhamnolipid production using agar plates containing methylene blue and cetyl trimethylammonium bromide. Biotechnol Lett. 2009;31(10):1583–8.

33. Siegmund I, Wagner F, Siegmund I, et al. New method for detecting rhamnolipids excreted byPseudomonas species during growth on mineral agar. Biotechnology Techniques. 1991;5(4):265–8.

34. Eric, Beck J, Evan, et al. Database resources of the National Center for Biotechnology Information in 2025. Nucleic Acids Research. 2025;53(D1):D20-D9.

35. Altschul SF, Gish W, Miller W, et al. Basic local alignment search tool. Journal of Molecular Biology. 1990;215(3):403–10.

36. Camacho C, Coulouris G, Avagyan V, et al. BLAST+: architecture and applications. BMC Bioinformatics. 2009;10(1):421.

37. Van den Bossche S, Vandeplassche E, Ostyn L, et al. Bacterial Interference With Lactate Dehydrogenase Assay Leads to an Underestimation of Cytotoxicity. Front Cell Infect Microbiol. 2020;10:494.

38. Tyanova S, Cox J. Perseus: A Bioinformatics Platform for Integrative Analysis of Proteom. Methods in Molecular Biology. 2018;1711:133–48.

39. Huang DW, Sherman BT, Lempicki RA, et al. Systematic and integrative analysis of large gene lists using DAVID bioinformatics resources. Nature Protocols. 2008;4(1):44–57.

40. Sherman BT, Hao M, Qiu J, et al. DAVID: a web server for functional enrichment analysis and functional annotation of gene lists (2021 update). Nucleic Acids Research. 2022;50(W1):W216–W21.

41. Winsor G, Griffiths E, Lo R, et al. Enhanced annotations and features for comparing thousands of Pseudomonas genomes in the Pseudomonas genome database. Nucleic acids research. 2016;44(D1):D646–53.

42. Stover C, Pham X, Erwin A, et al. Complete genome sequence of *Pseudomonas aeruginosa* PAO1, an opportunistic pathogen. Nature. 2000;406(6799):959–64.

43. Lewin GR, Kapur A, Cornforth DM, et al. Application of a quantitative framework to improve the accuracy of a bacterial infection model. Proceedings of the National Academy of Sciences. 2023;120(19):e2221542120.

44. Rossi E, Falcone M, Molin S, et al. High-resolution in situ transcriptomics of *Pseudomonas aeruginosa* unveils genotype independent patho-phenotypes in cystic fibrosis lungs. Nat Commun. 2018;9(1):3459.

45. Benjamini Y, Hochberg Y. Controlling the False Discovery Rate: A Practical and Powerful Approach to Multiple Testing. Journal of the Royal Statistical Society Series B (Methodological). 1995;57(1):289–300.

46. Team RC. R: A Language and Environment for Statistical Computing. Vienna, Austria: Foundation for Statistical Computing; 2024.

47. Wickham H. ggplot2: Elegant Graphics for Data Analysis. Springer-Verlag New York; 2016.

48. Guragain M, King MM, Williamson KS, et al. The *Pseudomonas aeruginosa* PAO1 Two-Component Regulator CarSR Regulates Calcium Homeostasis and Calcium-Induced Virulence Factor Production through Its Regulatory Targets CarO and CarP. Journal of Bacteriology. 2016;198(6):951–63.

49. Jude F, KöHler T, Branny P, et al. Posttranscriptional Control of Quorum-Sensing-Dependent Virulence Genes by DksA in *Pseudomonas aeruginosa*. Journal of Bacteriology. 2003;185(12):3558–66.

50. Min KB, Yoon SS. Transcriptome analysis reveals that the RNA polymerase–binding protein DksA1 has pleiotropic functions in *Pseudomonas aeruginosa*. Journal of Biological Chemistry. 2020;295(12):3851–64.

51. Min KB, Hwang W, Lee K-M, et al. Chemical inhibitors of the conserved bacterial transcriptional regulator DksA1 suppressed quorum sensing-mediated virulence of *Pseudomonas aeruginosa*. Journal of Biological Chemistry. 2021;296:100576.

52. Fortuna A, Bähre H, Visca P, et al. The two *Pseudomonas aeruginosa* DksA stringent response proteins are largely interchangeable at the whole transcriptome level and in the control of virulence-related traits. Environmental Microbiology. 2021;23(9):5487–504.

53. Fortuna A, Collalto D, Schiaffi V, et al. The *Pseudomonas aeruginosa* DksA1 protein is involved in H2O2 tolerance and within-macrophages survival and can be replaced by DksA2. Scientific Reports. 2022;12(1):10404.

54. Abergel C, Monchois V, Byrne D, et al. Structure and evolution of the Ivy protein family, unexpected lysozyme inhibitors in Gram-negative bacteria. Proceedings of the National Academy of Sciences. 2007;104(15):6394–9.

55. Clarke CA, Scheurwater EM, Clarke AJ. The Vertebrate Lysozyme Inhibitor Ivy Functions to Inhibit the Activity of Lytic Transglycosylase. Journal of Biological Chemistry. 2010;285(20):14843–7.

56. Spínola-Amilibia M, Davó-Siguero I, Ruiz FM, et al. The structure of VgrG1 from *Pseudomonas aeruginosa*, the needle tip of the bacterial type VI secretion system. Acta Crystallographica Section D Structural Biology. 2016;72(1):22–33.

57. Yang F, Gu J, Yang L, et al. Protective Efficacy of the Trivalent *Pseudomonas aeruginosa* Vaccine Candidate PcrV-OprI-Hcp1 in Murine Pneumonia and Burn Models. Scientific Reports. 2017;7(1):3957.

58. Milla CE, Chmiel JF, Accurso FJ, et al. Anti-PcrV antibody in cystic fibrosis: A novel approach targeting *Pseudomonas aeruginosa* airway infection. Pediatric Pulmonology. 2014;49(7):650–8.

59. Robb CS, Assmus M, Nano FE, et al. Structure of the T6SS lipoprotein TssJ1 from *Pseudomonas aeruginosa*. Acta Crystallographica Section F Structural Biology and Crystallization Communications. 2013;69(6):607–10.

60. Chen L, Zou Y, Kronfl AA, et al. Type VI secretion system of *Pseudomonas aeruginosa* is associated with biofilm formation but not environmental adaptation. MicrobiologyOpen. 2020;9(3):e991.

61. Tang Y, Romano FB, Breña M, et al. The *Pseudomonas aeruginosa* type III secretion translocator PopB assists the insertion of the PopD translocator into host cell membranes. Journal of Biological Chemistry. 2018;293(23):8982–93.

62. Damron FH, Napper J, Teter MA, et al. Lipotoxin F of *Pseudomonas aeruginosa* is an AlgU-dependent and alginate-independent outer membrane protein involved in resistance to oxidative stress and adhesion to A549 human lung epithelia. Microbiology. 2009;155(4):1028–38.

63. Firoved AM, Ornatowski W, Deretic V. Microarray Analysis Reveals Induction of Lipoprotein Genes in Mucoid *Pseudomonas aeruginosa*: Implications for Inflammation in Cystic Fibrosis. Infection and Immunity. 2004;72(9):5012–8.

64. Mudipalli Elavarasu S, K S. Rational design of an epitope-centric vaccine against *Pseudomonas aeruginosa* using pangenomic insights and immunoinformatics approach. Frontiers in Immunology. 2025;16:1617251.

65. Yang F, Gu J, Zou J, et al. PA0833 Is an OmpA C-Like Protein That Confers Protection Against *Pseudomonas aeruginosa* Infection. Frontiers in Microbiology. 2018;9:1062.

66. Zou J-T, Jing H-M, Yuan Y, et al. Pore-forming alpha-hemolysin efficiently improves the immunogenicity and protective efficacy of protein antigens. PLOS Pathogens. 2021;17(7):e1009752.

67. Fernández L, Breidenstein EBM, Song D, et al. Role of Intracellular Proteases in the Antibiotic Resistance, Motility, and Biofilm Formation of *Pseudomonas aeruginosa*. Antimicrobial Agents and Chemotherapy. 2012;56(2):1128–32.

68. Couto N, Schooling SR, Dutcher JR, et al. Proteome Profiles of Outer Membrane Vesicles and Extracellular Matrix of *Pseudomonas aeruginosa* Biofilms. Journal of Proteome Research. 2015;14(10):4207–22.

69. Toyofuku M, Roschitzki B, Riedel K, et al. Identification of Proteins Associated with the *Pseudomonas aeruginosa* Biofilm Extracellular Matrix. Journal of Proteome Research. 2012;11(10):4906–15.

70. Choi DS, Kim DK, Choi SJ, et al. Proteomic analysis of outer membrane vesicles derived from *Pseudomonas aeruginosa*. Proteomics. 2011;11(16):3424–9.

71. Ebner P, Götz F. Bacterial Excretion of Cytoplasmic Proteins (ECP): Occurrence, Mechanism, and Function. Trends in Microbiology. 2019;27(2):176–87.

72. Bjarnsholt T, Jensen PØ, Jakobsen TH, et al. Quorum Sensing and Virulence of *Pseudomonas aeruginosa* during Lung Infection of Cystic Fibrosis Patients. PLoS ONE. 2010;5(4):e10115.

73. Weimann A, Dinan AM, Ruis C, et al. Evolution and host-specific adaptation of *Pseudomonas aeruginosa*. Science. 2024;385(6704):eadi0908.

74. Blaby-Haas CE, Furman R, Rodionov DA, et al. Role of a Zn-independent DksA in Zn homeostasis and stringent response. Molecular Microbiology. 2011;79(3):700–15.

75. Furman R, Biswas T, Danhart EM, et al. DksA2, a zinc-independent structural analog of the transcription factor DksA. FEBS Letters. 2013;587(6):614–9.

76. Mastropasqua MC, Lamont I, Martin LW, et al. Efficient zinc uptake is critical for the ability of *Pseudomonas aeruginosa* to express virulence traits and colonize the human lung. Journal of Trace Elements in Medicine and Biology. 2018;48:74–80.

77. Colautti J, Kelly SD, Whitney JC. Specialized killing across the domains of life by the type VI secretion systems of *Pseudomonas aeruginosa*. Biochemical Journal. 2025;482(01):1–15.

78. Hauser AR. The type III secretion system of *Pseudomonas aeruginosa*: infection by injection. Nature Reviews Microbiology. 2009;7(9):654–65.

79. Galle M, Schotte P, Haegman M, et al. The *Pseudomonas aeruginosa* Type III secretion system plays a dual role in the regulation of caspase-1 mediated IL-1beta maturation. J Cell Mol Med. 2008;12(5A):1767–76.

80. Miao EA, Ernst RK, Dors M, et al. *Pseudomonas aeruginosa* activates caspase 1 through Ipaf. Proc Natl Acad Sci U S A. 2008;105(7):2562–7.

81. Sen-Kilic E, Huckaby AB, Damron FH, et al. *P. aeruginosa* type III and type VI secretion systems modulate early response gene expression in type II pneumocytes *in vitro*. Bmc Genomics. 2022;23(1):345.

82. Kolbe U, Yi B, Poth T, et al. Early Cytokine Induction Upon *Pseudomonas aeruginosa* Infection in Murine Precision Cut Lung Slices Depends on Sensing of Bacterial Viability. Front Immunol. 2020;11:598636.

83. Bomberger JM, Maceachran DP, Coutermarsh BA, et al. Long-distance delivery of bacterial virulence factors by *Pseudomonas aeruginosa* outer membrane vesicles. PLoS Pathog. 2009;5(4):e1000382.

84. Qin S, Xiao W, Zhou C, et al. *Pseudomonas aeruginosa*: pathogenesis, virulence factors, antibiotic resistance, interaction with host, technology advances and emerging therapeutics. Signal Transduct Target Ther. 2022;7(1):199.

85. Kim SI, Kim S, Kim E, et al. Secretion of Salmonella Pathogenicity Island 1-Encoded Type III Secretion System Effectors by Outer Membrane Vesicles in Salmonella enterica Serovar Typhimurium. Front Microbiol. 2018;9:2810.

86. Bai J, Kim SI, Ryu S, et al. Identification and characterization of outer membrane vesicle-associated proteins in Salmonella enterica serovar Typhimurium. Infect Immun. 2014;82(10):4001–10.

87. Sirisaengtaksin N, O’Donoghue EJ, Jabbari S, et al. Bacterial outer membrane vesicles provide an alternative pathway for trafficking of Escherichia coli O157 type III secreted effectors to epithelial cells. mSphere. 2023;8(6):e0052023.

88. Armstrong DA, Lee MK, Hazlett HF, et al. Extracellular Vesicles from *Pseudomonas aeruginosa* Suppress MHC-Related Molecules in Human Lung Macrophages. ImmunoHorizons. 2020;4(8):508-19.

